# Facile generation of biepitopic antibodies with intrinsic agonism for activating receptors in the tumor necrosis factor superfamily

**DOI:** 10.1101/2023.12.11.571146

**Authors:** Harkamal S. Jhajj, John S. Schardt, Namir Khalasawi, Emily L. Yao, Timon S. Lwo, Na-Young Kwon, Ryen L O’Meara, Alec A. Desai, Peter M. Tessier

## Abstract

Agonist antibodies that activate cellular receptors are being pursued for therapeutic applications ranging from neurodegenerative diseases to cancer. For the tumor necrosis factor (TNF) receptor superfamily, higher-order clustering of three or more receptors is key to their potent activation. This can be achieved using antibodies that recognize two unique epitopes on the same receptor and mediate receptor superclustering. However, identifying compatible pairs of antibodies to generate biepitopic antibodies (also known as biparatopic antibodies) for activating TNF receptors typically requires animal immunization and is a laborious and unpredictable process. Here, we report a simple method for systematically identifying biepitopic antibodies that potently activate TNF receptors without the need for additional animal immunization. Our approach uses off-the-shelf, receptor-specific IgG antibodies, which lack intrinsic (Fc-gamma receptor-independent) agonist activity, to first block their corresponding epitopes. Next, we perform selections for single-chain antibodies from human nonimmune libraries that bind accessible epitopes on the same ectodomains using yeast surface display and fluorescence-activated cell sorting. The selected single-chain antibodies are finally fused to the light chains of IgGs to generate human tetravalent antibodies that engage two different receptor epitopes and mediate potent receptor activation. We highlight the broad utility of this approach by converting several existing clinical-stage antibodies against TNF receptors, including ivuxolimab and pogalizumab against OX40 and utomilumab against CD137, into biepitopic antibodies with highly potent agonist activity. We expect that this widely accessible methodology can be used to systematically generate biepitopic antibodies for activating other receptors in the TNF receptor superfamily and many other receptors whose activation is dependent on strong receptor clustering.

## Introduction

Antibodies that target immune checkpoints are an important class of emerging therapeutics due to their ability to regulate immune cell function (Waldman et al., 2020). In the context of cancer, the tumor microenvironment is often plagued by immunosuppression, which can be reversed by targeting co-inhibitory and co-stimulatory receptors on T cells (Lu et al., 2020; Zahavi and Weiner, 2020). Tumor cells have been shown to hijack these immune checkpoints to evade clearance by the host immune response (Han et al., 2020). Specifically, monoclonal antibodies that block the immunosuppressive interactions on co-inhibitory receptors have shown potent anti-tumor responses. Encouragingly, numerous immunotherapies have been developed for these inhibitory checkpoints and many have received FDA approval for treating a variety of cancers, including those against PD-1 (*e.g.*, pembrolizumab, nivolumab, and cemiplimab), PD-L1 (*e.g.*, atezolizumab, durvalumab, and avelumab) and CTLA-4 (*e.g.*, ipilimumab) (Lee et al., 2019). While these therapies have demonstrated impressive efficacies, the benefits are limited to only a subset of patients and there remains an urgent clinical need for alternative or complementary immunotherapy approaches to mediate potent anti-tumor responses (Callahan et al., 2016; Chowdhury et al., 2018).

The development of agonist antibodies that activate co-stimulatory checkpoints, such as the tumor necrosis factor (TNF) receptor superfamily (*e.g.,* OX40, CD137, CD40), has garnered intense interest. This is evident by the myriad of ongoing, early-stage clinical trials evaluating TNF receptor agonist monotherapies and combination immunotherapies in oncologic applications (Hahn et al., 2017; Jhajj et al., 2023; Mayes et al., 2018; Schardt et al., 2022). Despite their promise, their clinical translation has been hindered by safety and efficacy concerns (Sedger and McDermott, 2014). In particular, OX40 and CD137 agonist antibodies (*i.e.,* bivalent IgGs) have been limited by poor-to-modest clinical activity, which is not surprising given that complete activation of OX40 and CD137 (like other TNF receptor superfamily members) requires trimerization and subsequent higher-order receptor superclustering (Chin et al., 2018; Melero et al., 2013). Bivalent IgGs typically rely on engagement with Fcγ receptors (FcγRs) on antigen-presenting cells to achieve such higher-order receptor clustering (Kim and Ashkenazi, 2013; Wilson et al., 2011). However, FcγR expression is observed to vary greatly between different immune cells and is difficult to predict and control *in vivo* (Furness et al., 2014; Kim and Ashkenazi, 2013). Overall, novel strategies to activate immune co-stimulatory receptors systematically and reliably in an FcγR-independent manner are anticipated to improve the therapeutic limitations associated with modest clinical activity.

Recently, biepitopic antibodies targeting two non-overlapping receptor regions have emerged as a robust strategy to activate immune co-stimulatory receptors via the induction of extensive daisy-chain-like receptor superclusters (Yang et al., 2019). Encouragingly, potent agonist activity has been observed *in vivo* for tetravalent, biepitopic OX40 antibodies lacking affinity for FcγRs. Conventional methods of generating such antibodies generally involve four key steps: i) immunizing animals; ii) isolating individual antibodies; iii) epitope binning antibodies to identify pairs with unique epitopes; and iv) combining pairs of antibodies into non-conventional (non-IgG) formats that present two unique antigen-binding sites in a single antibody with an overall valency of three or more (Abdiche et al., 2012, 2009; Anderson et al., 2017).

Despite previous successes using this approach (Abdiche et al., 2014; Sivasubramanian et al., 2017), it suffers from four main problems. First, animal immunization and subsequent monoclonal antibody identification are slow and unpredictable processes. Second, it is challenging to reliably obtain sufficient epitope coverage, especially due to immunodominant epitopes. Third, the resulting antibodies typically need to be humanized for therapeutic applications. Even in cases where animals with humanized immune systems are used, the benefits of not needing to humanize are often diminished by reduced antibody diversity and epitope coverage. Fourth, the process of combining pairs of antibodies in non-native antibody formats – such as reformatting antibodies discovered as IgGs into single-chain antibodies (e.g., scFvs) to be fused to other IgGs – can result in partial or complete loss of binding of the single-chain antibodies.

Therefore, we sought to develop an approach that addresses each of these challenges and greatly simplifies the generation of biepitopic antibodies. First, we eliminate the need for additional animal immunization by using existing, off-the-shelf IgGs specific for the target receptor. While such antibodies are typically generated by immunization, there are many existing IgGs specific for most receptors of therapeutic interest and the key problem is converting them into biepitopic antibodies in a simple and systematic manner. Second, we perform *in vitro* selections of single-chain antibodies after blocking the receptor epitope of the existing IgG, which results in simple isolation of pairs of antibodies with unique epitopes. Third, the *in vitro* antibody selections are performed using human nonimmune libraries, which are the same libraries for every antigen and do not require humanization. Fourth, we perform *in vitro* selections for single-chain antibodies in the final antibody format that will be used to generate biepitopic antibodies, such as the IgG-single chain antibody format used in this work, which greatly reduces the risk of loss of binding due to reformatting (i.e., conversion of IgGs into scFvs). Herein, we demonstrate the utility of this approach using existing clinical-stage antibodies specific for multiple receptors in the TNF receptor superfamily that lack intrinsic (FcγR-independent) agonist activity and demonstrate how they can be systematically converted into biepitopic antibodies with potent activity to activate human T cells.

## Results

### Competition-based selection of OX40 single-chain antibodies

To develop biepitopic OX40 antibodies, we sought to identify single-chain antibodies (scFvs) to pair with a clinical-stage OX40 agonist antibody that lacks intrinsic agonist activity, namely 11D4 (also known as ivuxolimab, PF-04518600, PF-8600, and B110) (Diab et al., 2022). To establish proof-of-principle results for our competition-based discovery approach, we first evaluated the expression and antigen-binding activity of 11D4 as a single-chain Fab (scFab) fragment on the surface of yeast (**Figure S1**). Encouragingly, the 11D4 scFab expressed well and bound the OX40 extracellular domain. Moreover, we confirmed that pre-incubation of OX40 with 11D4 IgG effectively blocked binding to the yeast-displayed 11D4 scFab. This demonstrates our ability to specifically block the 11D4 epitope on OX40, which we aim to use for selecting antibodies that recognize non-overlapping OX40 epitopes.

Next, we performed a series of *in vitro* selections to identify scFvs that recognize unique OX40 epitopes from a human nonimmune library (Feldhaus et al., 2003; **Figure 1**). First, the library was enriched for OX40 binding via an initial round of magnetic-activated cell sorting (MACS) against bivalent OX40-Fc. The choice of using bivalent OX40 is because the initial step of enriching antibody binders from nonimmune libraires is typically more effective using multivalent antigens. Following one round of MACS, the yeast population was propagated and evaluated via fluorescence-activated cells sorting (FACS) for binding to biotinylated monovalent OX40. The decision to use monovalent antigen at the FACS stage of library sorting is because this typically enables better selection of the highest affinity antibodies. Significant enrichment of the antigen-binding population was observed after each sort (FACS rounds 1-3), as shown in **Figure 1**.

**Figure 1.**
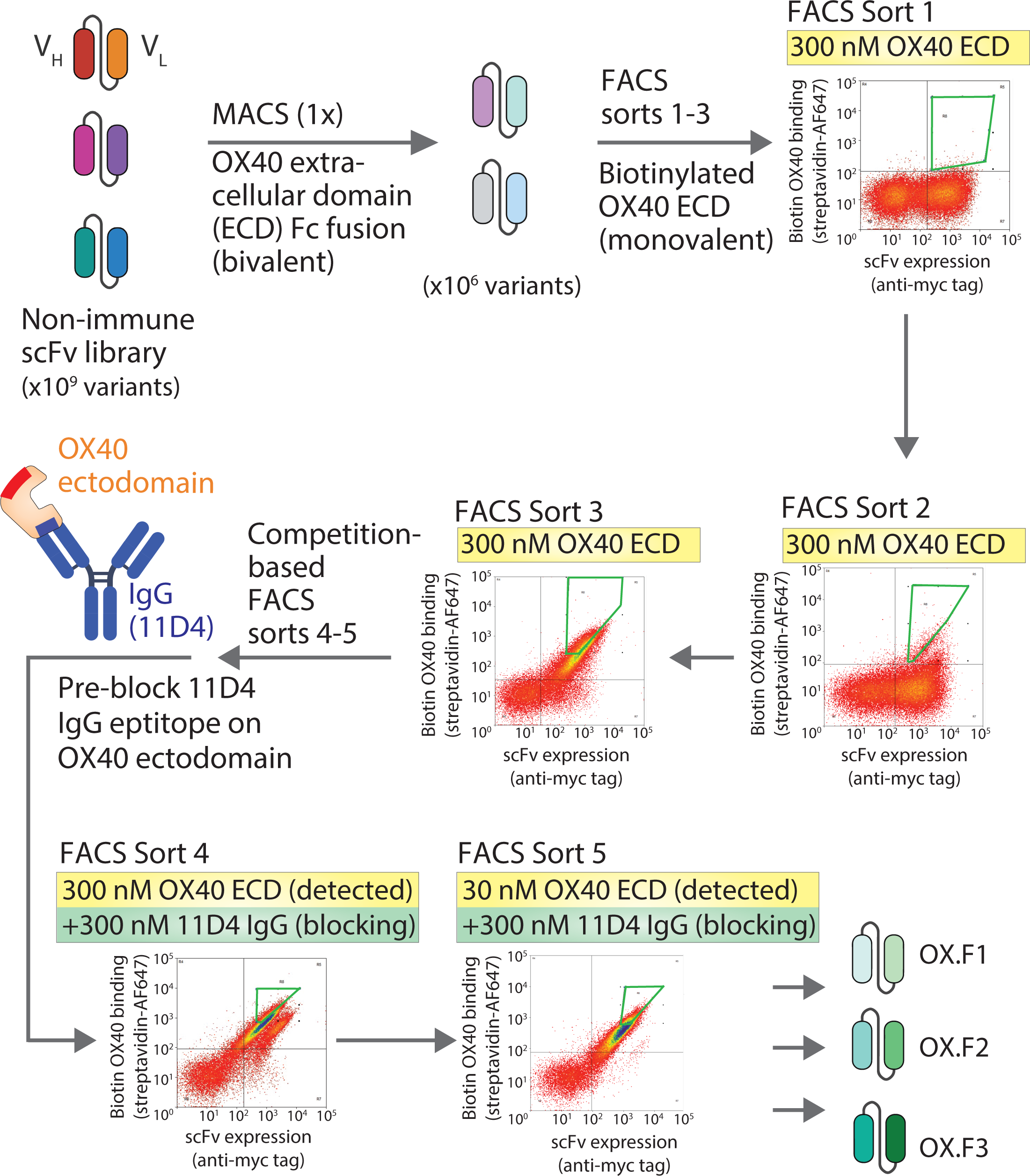
Overview of approach for isolating single-chain antibodies from human libraries with unique receptor (OX40) epitopes relative to a clinical-stage OX40 antibody. A human single-chain (scFv) library displayed on yeast was enriched for binding to the human OX40 ectodomain via magnetic-activated cell sorting (MACS, one sort) and fluores-cence-activated cell sorting (FACS, three sorts). Next, a clinical-stage OX40 IgG (11D4, also known as ivuxolimab) was used to block its OX40 epitope on the soluble OX40 ectodomain, and the library was further enriched against the OX40/IgG complex to identify scFvs with unique OX40 epitopes. Three unique scFvs (OX.F1, OX.F2, and OX.F3) were discovered using this approach. The figure was made using BioRender.

We next sought to identify scFvs that engaged non-overlapping OX40 epitopes relative to the 11D4 epitope. We applied a competition-based strategy by screening for antigen binding in the presence of soluble 11D4 IgG. Consecutive rounds of screening were conducted, first at an equimolar concentration of OX40:11D4 IgG followed by a 10-fold molar excess of 11D4 IgG relative to OX40. Encouragingly, we retained a clear binding population after pre-blocking the 11D4 epitope on OX40 (**Figure 1**; FACS sort 4) and subsequently observed significant library enrichment (FACS sort 5), suggesting an enrichment for antibodies with non-overlapping epitopes had been achieved. Finally, we sequenced antibodies from the enriched libraries and identified three unique variants (OX.F1, OX.F2, and OX.F3) for continued evaluation (**Figure S2**).

### Selected single-chain antibodies recognize unique epitopes relative to existing OX40 IgG

To evaluate the OX40 binding of the identified scFvs, affinity analysis on yeast was conducted in the presence or absence of 11D4 IgG pre-blocking (**Figure 2A**). Encouragingly, each of the three scFvs (OX.F1, OX.F2, and OX.F3) retained binding to OX40 in the presence and absence of 11D4 IgG, suggesting that the scFvs engage distinct epitopes (**Figures 2C** and **S3**). The scFvs exhibited stronger OX40 binding after 11D4 IgG pre-blocking compared to the control without pre-blocking. Importantly, we confirmed that the three scFvs did not bind 11D4 IgG (**Figure S4**). The increased OX40 binding for the three scFvs may be due to the increased avidity of the monovalent OX40 ectodomain, which has the potential to become bivalent after binding to the bivalent 11D4 IgG. As expected, 11D4 pre-blocking significantly reduced 11D4 scFab binding to OX40, confirming that antibodies with overlapping epitopes reduce OX40 binding (**Figure 2B**). Overall, these results demonstrate that the selected single-chain antibodies engage unique OX40 epitopes compared to 11D4 IgG.

**Figure 2.**
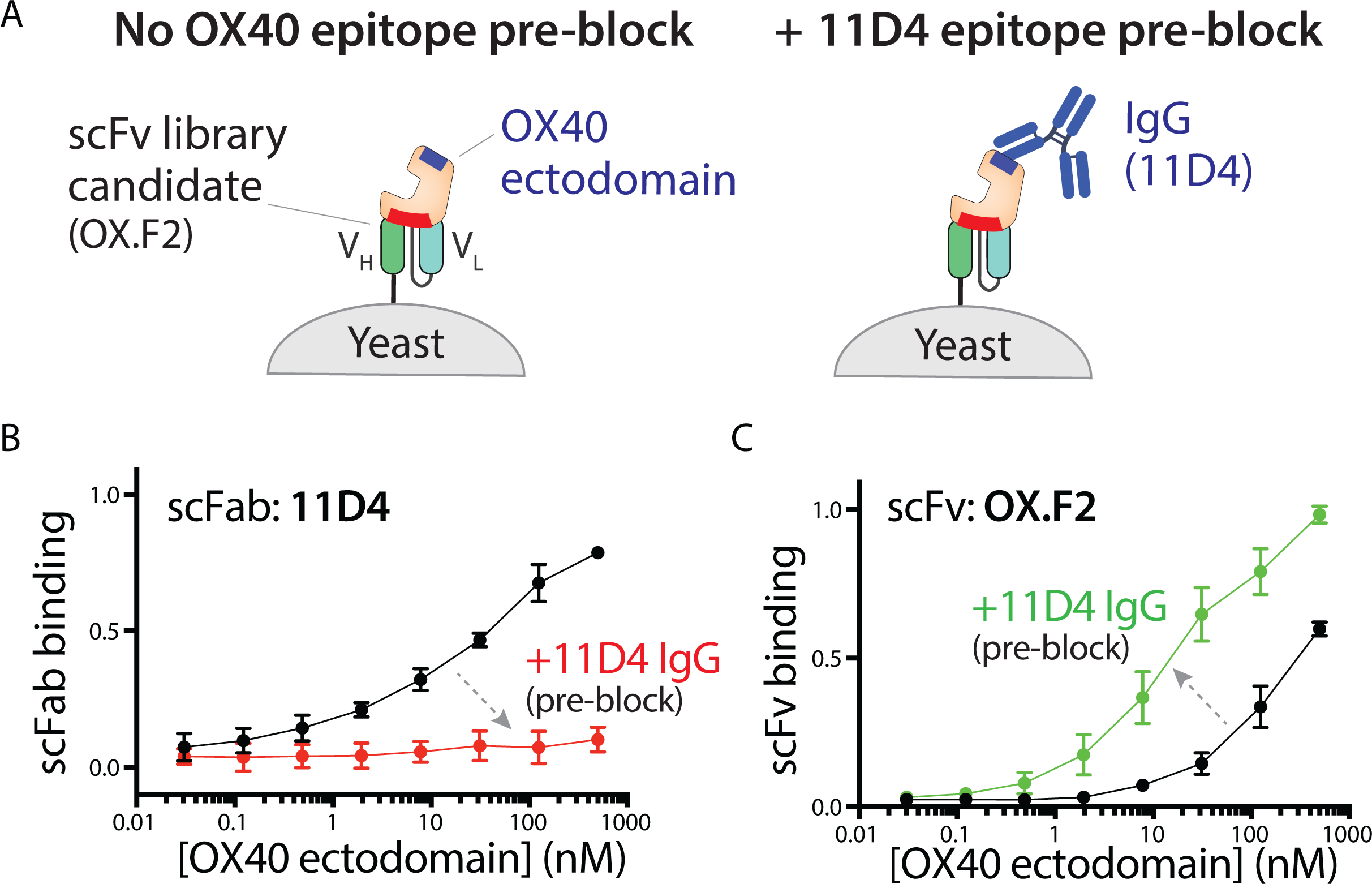
Selected single-chain antibody possesses unique OX40 epitope relative to 11D4 IgG. (A) The candidate single-chain antibody (OX.F2) was displayed on yeast and evaluated for binding to monovalent OX40 ectodomain when the 11D4 epitope is pre-blocked. (B) The wild-type 11D4 single-chain Fab (scFab) on yeast loses binding to OX40 when the ecto-domain is pre-incubated with 11D4 IgG, as expected. (C) Selected OX40 scFv (OX.F2) shows strong binding to OX40 in the presence and absence of 11D4 IgG pre-blocking. The results are averages of three independent experiments and the error bars are standard deviations.

Next, we investigated the binding of the selected scFvs as Fc-fusion proteins to OX40 on HEK cells to ensure that the clones bound the native OX40 receptor in addition to the recombinant extracellular domain. Encouragingly, the scFv-Fc fusion proteins showed dose-dependent binding to the OX40 receptor (**Figure S5**). Moreover, the OX.F2 antibody displayed higher apparent affinity than the OX.F1 and OX.F3 variants. Collectively, these results indicate that the selected scFvs bind to unique OX40 epitopes relative to 11D4 IgG and recognize the native cell-surface OX40 receptor, which motivates combining them into biepitopic antibodies and evaluating them as potent OX40 receptor agonists.

### Biepitopic OX40 antibodies show potent, Fc**γ**R-independent bioactivities in primary human CD4^+^ T cells

We next generated biepitopic antibodies – without any molecular reformatting of the 11D4 IgG or scFvs – by simply fusing the scFvs to the C-termini of the light chains of 11D4 IgG (Orcutt et al., 2010; **Figure 3A**). This approach is different than other reformatting methods in which IgGs are reformatted as scFvs or vice versa, as the latter approaches involve adding or removing peptide linkers between the variable regions and, in some cases, reduce binding affinity and/or stability due to changes in the Fv structure (Brinkmann and Kontermann, 2017; Tiller and Tessier, 2015). The resulting tetravalent antibodies are referred to as IgG-scFv antibodies. They expressed well and their purification yields were relatively high (12-17 mg/L for 11D4-OX.F2 and 11D4 IgG-scFv relative to 38±16 mg/L for 11D4 IgG), and they were isolated at high purity (94-99% monomer after two-step purification; **Figures S6** and **S7**). The biepitopic 11D4-OX.F2 antibody displayed generally similar binding profiles to OX40 HEK293 cells as the 11D4 IgG (**Figure S5**). The ability of the biepitopic IgG-scFv antibodies to promote receptor activation was first evaluated using engineered OX40 Jurkat cells that report intracellular signaling (NF-κB activation) due to receptor activation via luciferase expression (**Figure S8**). In the absence of FcγR-mediated crosslinking, the tetravalent 11D4-OX.F2 antibody induced strong NF-κB activity in a dose-dependent manner. In comparison, the bivalent 11D4 IgG was unable to induce receptor activation due to the lack of FcγR-mediated crosslinking.

**Figure 3.**
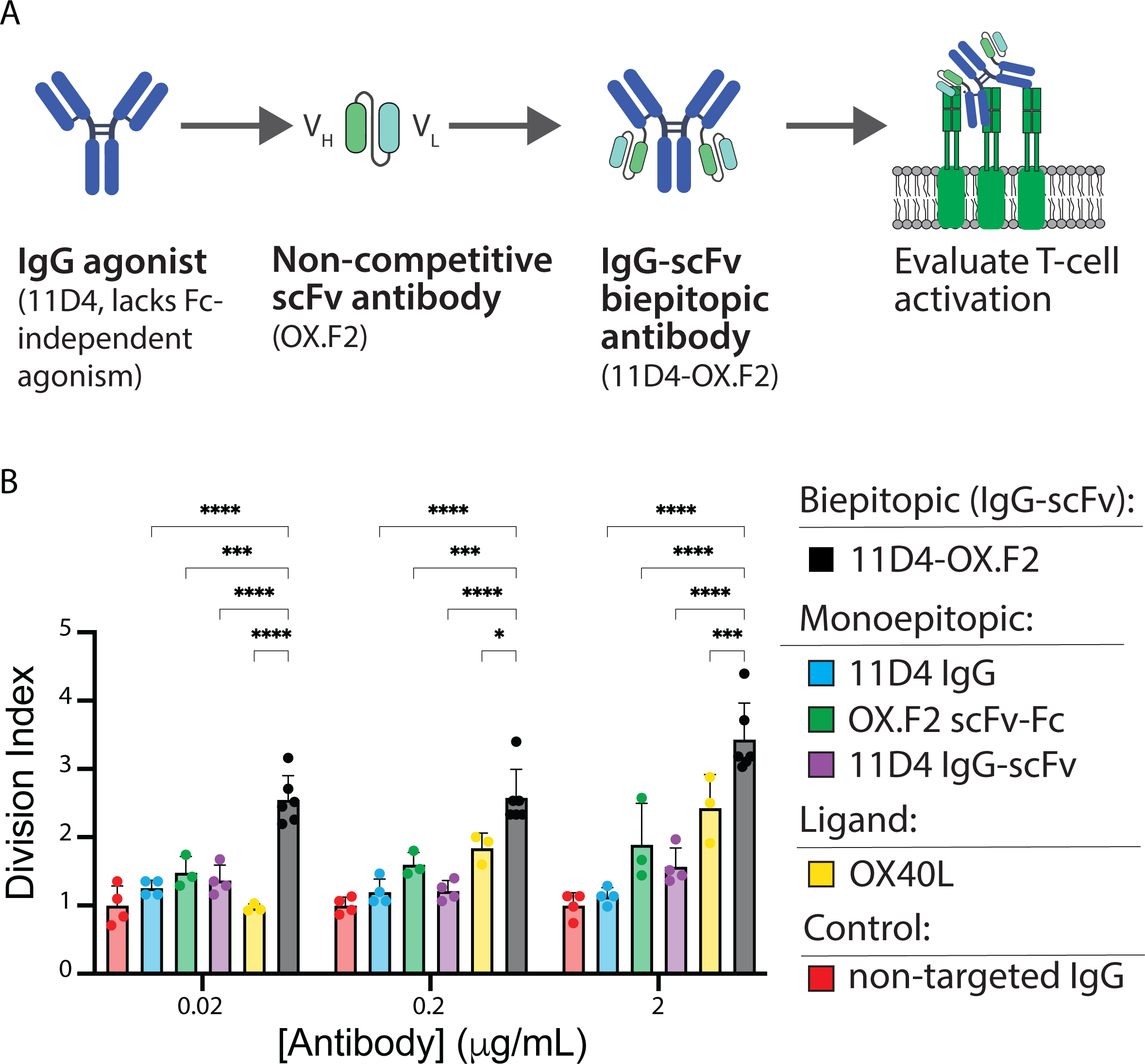
Biepitopic OX40 antibody induces strong human CD4^+^ T cell activation in an FcγR-independent manner. (A) Schematic illustration of the approach used to convert an existing clinical-stage antibody into a potent OX40 antibody with intrinsic agonist activity. (B) The biepitopic antibody induced strong human CD4^+^ T cell proliferation. Division index was calculated using the FlowJo Proliferation platform to evaluate proliferation at day 6. The results are averages of three independent experiments and the error bars are standard deviations. In (B), the biepitopic (IgG-scFv) antibodies displayed higher levels of proliferation than the monoepitopic antibodies and OX40 ligand at each concentration [*p*-value <0.05 (*), <0.01 (**), <0.001 (***), and <0.0001 (****)].

Next, we sought to examine the relative activities of the biepitopic IgG-scFv antibodies for their ability to induce human CD4^+^ T cell proliferation (**Figure 3A**). To evaluate their dose-dependent responses, CD4^+^ T cells were incubated with a broad range of concentrations of OX40 antibodies (0.02-2 μg/mL) along with anti-CD3/CD28 antibodies (primary/secondary T cell activators; **Figures 3B** and **S9**). The division index – the average number of cell divisions for a T cell in the original population – was determined, which demonstrated that the biepitopic IgG-scFv antibody (11D4-OX.F2) induced the highest and most potent levels of T cell proliferation. In contrast, the monoepitopic, tetravalent antibody (11D4 IgG-scFv) resulted in little T cell proliferation. Moreover, 11D4 IgG and OX.F2 scFv-Fc, which are the antibody components of the 11D4-OX.F2 tetravalent antibody, also induced little T cell proliferation, as expected given their lack of intrinsic receptor agonism due to their bivalent natures. Notably, the 11D4-OX.F2 antibody was also more active than the OX40 ligand, highlighting the potent nature of the biepitopic antibody. These data demonstrate the importance of multivalent antibodies in promoting strong receptor activation and improving agonist activity compared to the bivalent antibodies.

### Additional clinical-stage OX40 IgGs engineered as biepitopic antibodies activate primary human CD4^+^ T cells

Next, we investigated the generality of our findings – that biepitopic IgG-scFv antibodies induce superior and FcγR-independent activities relative to their monoepitopic counterparts – by identifying other clinical-stage OX40 antibodies to pair with OX.F2 and evaluating them for bioactivity in primary T cell models (**Figure 4A**). Toward this goal, we first conducted competitive binding analysis to compare the epitope of other clinical-stage OX40 IgGs (pogalizumab, 18D8, and telazorlimab) relative to that of 11D4 and OX.F2. All three clinical-stage antibodies did not compete with 11D4 for OX40 binding (**Figure 4B**), revealing that they possess unique epitopes. Interestingly, pogalizumab and 18D8 engage a unique epitope relative to OX.F2, while telazorlimab engages an overlapping epitope (**Figure 4C**). These findings were further corroborated by a second competitive binding assay (**Figure S10**). All together, these results demonstrate that OX.F2 engages a unique OX40 epitope relative to the additional clinical-stage OX40 IgGs (pogalizumab and 18D8) and motivates combining them as IgG-scFv biepitopic antibodies and evaluating their function to assess the generality of our approach.

**Figure 4.**
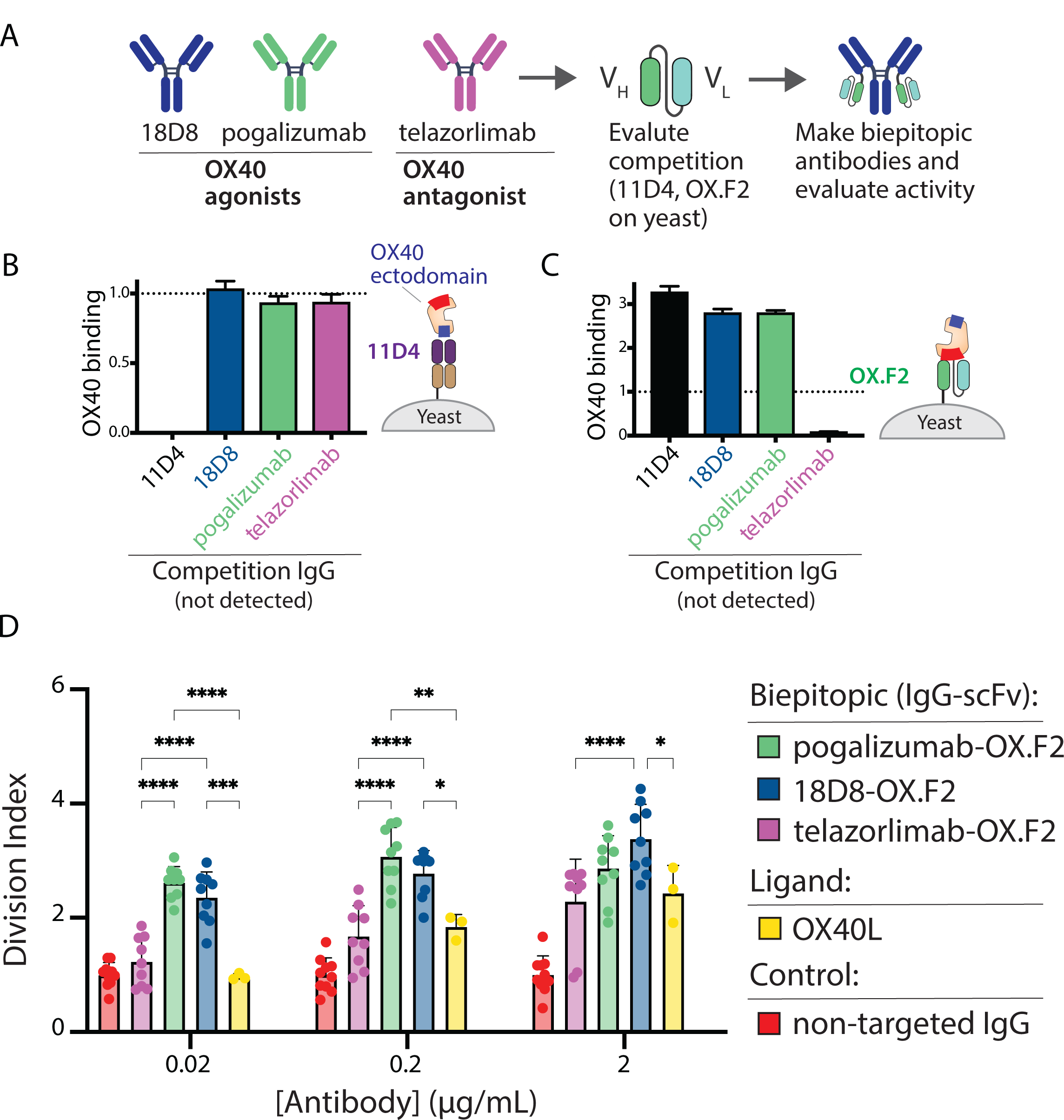
Generalization of the biepitopic antibody approach to additional OX40 clinical-stage antibodies results in potent human CD4^+^ T cell activation. (A) Schematic of the process of competition-based analysis of antibody epitopes for clinical-stage OX40 agonists and antagonists, the resulting combination of IgGs with the OX.F2 scFv into biepitopic formats, and their evaluation for T cell activation. (B-C) Competition-based analysis of (B) 11D4 and (C) OX.F2 single-chain antibodies displayed on yeast binding to OX40 after pre-blocking OX40 with a clinical-stage OX40 antibody. (D) The biepitopic antibodies induced strong human CD4^+^ T cell proliferation. Division index was calculated using the FlowJo Proliferation platform to evaluate proliferation at day 6. The results are averages of three independent experiments and the error bars are standard deviations. 18D8-OX.F2 displayed higher levels of proliferation than telazorlimab-OX.F2 and OX40 ligand at each concentration, while pogalizumab-OX.F2 displayed higher levels of proliferation than telazorlimab-OX.F2 and OX40 ligand at 0.02 and 0.2 μg/mL [*p*-value <0.05 (*), <0.01 (**), <0.001 (***), and <0.0001 (****)].

Therefore, we evaluated the ability of the OX.F2 scFv paired with the panel of clinical-stage OX40 IgGs to activate OX40 on human CD4^+^ T cells (**Figure 4D**). We first generated the tetravalent antibodies, which were purified at relatively high yield (9-19 mg/L for the tetravalent antibodies relative to 20-27 mg/L for the IgGs) and purity (98-99% monomer after two-step purification; **Figures S6** and **S7**). We observed that the two biepitopic antibodies, pogalizumab-OX.F2 and 18D8-OX.F2, mediated strong CD4^+^ T cell proliferation in a similar manner to 11D4-OX.F2 (**Figure 4D**). Interestingly, telazorlimab-OX.F2 elicited weaker T cell proliferation, which was expected based on the competitive binding analysis that revealed telazorlimab and OX.F2 recognize overlapping epitopes (**Figure 4C**). Finally, we observed that the biepitopic antibodies more potently activated OX40 than its cognate ligand (**Figure 4D**). Overall, these data demonstrate that the OX.F2 single-chain antibody can be paired with existing clinical-stage OX40 IgGs – without any molecular reformatting and only requiring fusing two antibodies together – to generate biepitopic antibodies with strong ability to activate human T cells.

### Competition-based discovery platform can be generalized to other TNF receptors

To evaluate the broad applicability of the antibody screening platform, we sought to discover scFvs that could be paired with a clinical-stage CD137 antibody that lacks intrinsic agonist activity, namely utomilumab (Chin et al., 2018). To do this, we used the same discovery approach we established for OX40 (Kelly et al., 2018; **Figure 5A**). We first used MACS to enrich the antibody library against the CD137 ectodomain and then further enriched the library used FACS (sorts 1-4). Next, competition-based screening was performed by pre-blocking biotinylated CD137 with utomilumab IgG and selecting scFvs that maintained CD137 binding (sorts 5-6). This included FACS sorts that used equimolar concentrations of CD137 and utomilumab IgG (sort 5) followed by sorts that used excess utomilumab IgG relative to CD137 (sort 6). Finally, the enriched libraries were sequenced and one scFv was identified, namely CD.K2 (**Figure S11**).

**Figure 5.**
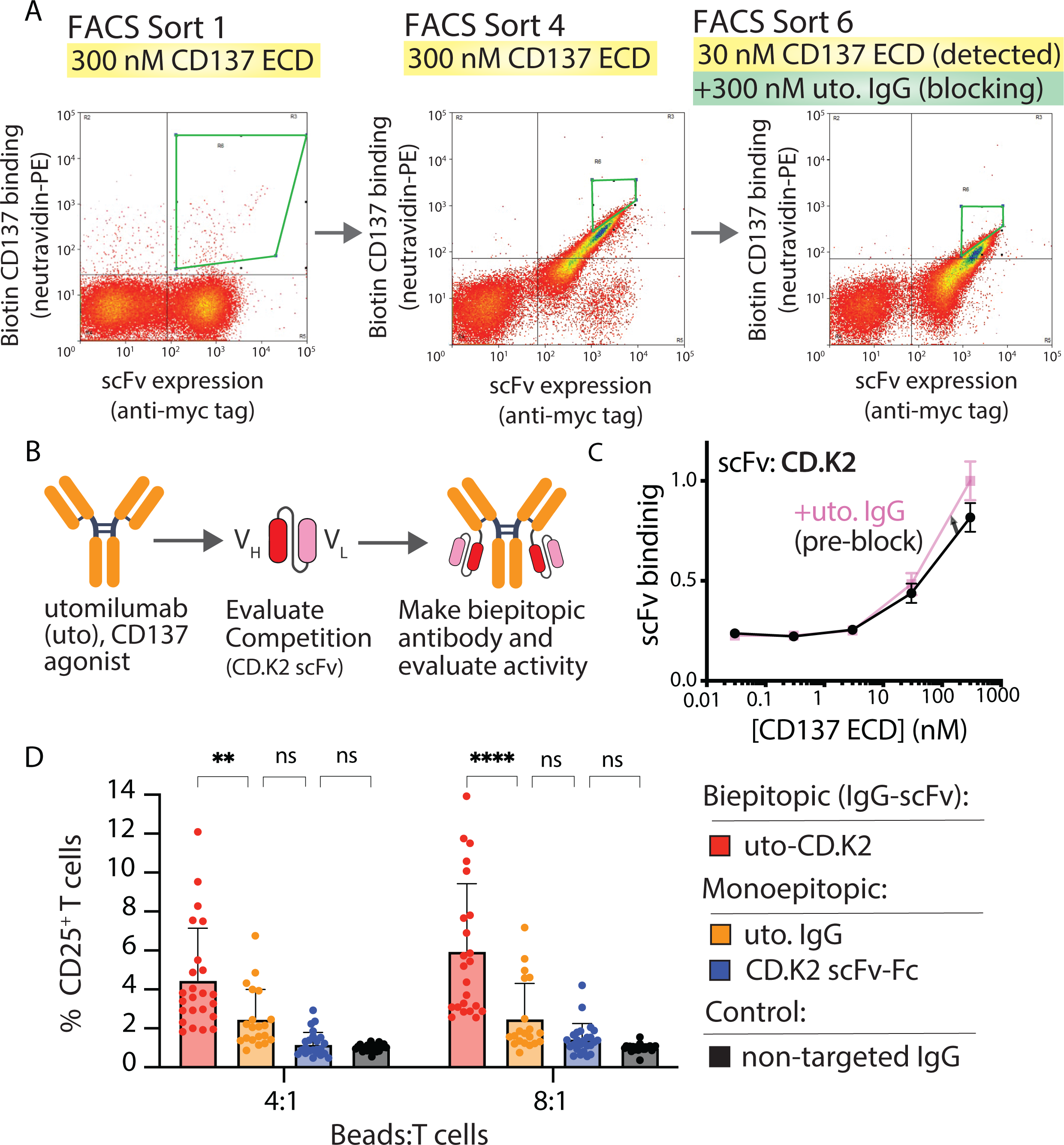
Generalization of the biepitopic antibody approach to an additional TNF receptor (CD137) results in potent human CD8^+^ T cell activation. (A) A human scFv library was enriched against the human CD137 ectodomain via MACS (one sort) and FACS (three sorts). Next, the library was enriched for CD137 binding in the presence of a clinical-stage CD137 IgG (utomilumab, uto.; two sorts). (B) Schematic of the approach used to convert utomilumab into a biepitopic CD137 antibody with intrinsic agonist activity. (C) The isolated CD137 single-chain antibody (CD.K2) displayed on yeast bound CD137 ectodomain in the absence and presence of uto IgG. (D) The biepitopic antibody (utomilumab-CD.K2; uto-CD.K2) induced strong human CD8^+^ T cell proliferation, as judged by CD25 expression, relative to the monoepitopic antibodies and a control IgG. The results are averages of five independent experiments and the error bars are standard errors [*p*-value <0.01 (**) and <0.0001 (****)].

The binding and functional analysis of CD.K2 scFv was performed in multiple steps (**Figure 5B**). First, we evaluated the binding of CD.K2 to recombinant CD137 in the presence and absence of utomilumab IgG pre-blocking and did not observe competition (**Figure 5C**), suggesting that the two antibodies recognize unique CD137 epitopes. As a positive control, we confirmed that the utomilumab scFv lost binding in the presence of utomilumab IgG pre-blocking (**Figure S12**). We also generated CD.K2 as a soluble Fc-fusion protein and confirmed it bound to CD137 on HEK293 cells (**Figure S13**). Finally, we generated an IgG-scFv antibody using the CD.K2 scFv (**Figures 5B, S14** and **S15**) and tested its ability to activate human primary CD8^+^ T cells (**Figures 5D** and **S16**). Encouragingly, the biepitopic antibody (uto-CD.K2) displayed strong T cell activation, as judged by increased CD25 levels, while the monoepitopic antibodies (uto. IgG and CD.K2 scFv-Fc) did not. Collectively, these results demonstrate that the competition-based antibody screening platform can be generalized to other members of the TNF receptor superfamily and used to greatly simplify the development of potent T cell agonist antibodies.

## Discussion

To generate potent agonists of OX40 and CD137, we developed a competition-based screening platform that is systematic, high-throughput, and easy to use. Our approach has several advantages relative to conventional methods that use a combination of animal immunization, primary antibody identification, and pair-wise screening of complementary antibodies based on epitope specificity (Abdiche et al., 2014, 2012, 2009; Anderson et al., 2017). The latter process is especially challenging because, at least in some cases, most isolated antibodies target relatively few immunodominant epitopes. For example, one study sought to discover antibodies against unique epitopes of a target protein (mesothelin) using hybridoma technology (Zhang et al., 2015). Among the 7,680 hybridomas, only 232 clones were specific for the target antigen and 96% (223 clones) shared the same epitope. Even conventional *in vitro* selection strategies are typically challenging and time consuming due to the need for several steps, including generating immune libraries, selecting primary antibodies via display technologies, reformatting the selected antibodies as soluble antibodies, performing selections for additional antibodies with orthogonal epitopes, and finally reformatting into biepitopic antibodies (Bogen et al., 2020). To overcome these challenges, our approach uses off-the-shelf IgGs that can be readily converted into biepitopic antibodies. We anticipate that this screening platform can be used for discovering a variety of novel biepitopic antibodies for a broad range of applications.

Our findings also highlight the importance of engineering multivalent antibodies to enhance OX40 and CD137 receptor activation, which is consistent with previous reports of using multivalency to improve agonist activity (Sadeghnezhad et al., 2019; Tigue et al., 2017; Yang et al., 2019). In our work, we designed tetravalent antibodies that exhibited improved NF-κB activation and T cell proliferation. These results are consistent with previous studies for the OX40 receptor where tetravalent antibody constructs, such as those in the dual-variable-domain (DVD) format, were shown to impart superior agonism due to their ability to promote high-order receptor clustering (Yang et al., 2019). Moreover, the DVD antibodies mediated significantly improved NF-κB activation compared to their bivalent counterparts. One explanation for enhanced agonist activity with higher valency is the requirement for trimerization of several receptors in the TNF receptor superfamily (Kucka and Wajant, 2021), which can be achieved using antibodies possessing at least three antigen-binding sites. To test this hypothesis, investigators engineered trivalent and tetravalent nanobody formats against death receptor 5 (DR-5). They found that these formats strongly decreased tumor viability and displayed an increase in apoptotic response compared to the corresponding bivalent nanobody (Sadeghnezhad et al., 2019). Antibodies with even higher valency, such as hexavalent antibodies, have also been explored to improve TNF receptor activation. In the case of the glucocorticoid-induced TNF receptor-related protein, a hexameric Fc-fusion protein (MEDI1873) exhibited significantly improved T cell proliferation relative to its bivalent IgG counterpart. These findings were further corroborated in a primate model where MEDI1873 showed enhanced T cell proliferation and elevated levels of IgG circulating antibodies, suggesting both strong cellular and humoral immune responses (Tigue et al., 2017). Beyond TNF receptors, more work is needed to evaluate the potential of multivalent antibodies for strongly activating other T cell receptors.

In addition to valency, our results also highlight significant impacts of engaging multiple receptor epitopes, which builds on previous findings related to strong activation of multiple TNF receptors (Bogen et al., 2020; Overdijk et al., 2020; Yang et al., 2019). Overall, biepitopic antibodies with superior bioactivities relative to their monoepitopic counterparts have been generated in a variety of molecular formats, including as tetravalent mAb-scFvs (as used in this work), tetravalent DVDs (Yang et al., 2019), bivalent IgGs (Bogen et al., 2020), and Fc-engineered constructs that non-covalently trimerize upon antigen binding (Overdijk et al., 2020). A previous report has shown that biepitopic antibodies in a bivalent IgG format induce receptor clustering of soluble and cell-bound epidermal growth factor receptors (EGFR) (Bogen et al., 2020). The authors hypothesized that the binding kinetics of biepitopic antibodies are favorable for receptor clustering due to a mixture of low and high-affinity Fab arms that lead to continuous binding and releasing effects to induce receptor superclusters. This phenomenon is even more amplified by tetravalent, biepitopic antibodies where multiple receptor clusters can be brought together to induce greater receptor activation.

The superior activity of biepitopic antibodies can also be explained by the theoretical extensive daisy-chain-like receptor superclustering made possible via intermolecular receptor engagement as opposed to terminating, intramolecular receptor engagement (Li et al., 2016; Wang et al., 2019; Yang et al., 2019). Molecular geometry, linker length, and linker rigidity are expected to influence such molecular interactions, and by extension, molecular engineering can be employed to bias toward desired molecular interactions. Biochemical methods, such as size-exclusion chromatography, and cell-based methods, such as fluorescence microscopy, have been established for characterizing the influence of antibody molecular format on receptor clustering, which may be useful for rapidly identifying the most effective formats and epitopes that maximize receptor activation (Li et al., 2016; Yang et al., 2019).

Conversely, biepitopic antibodies have also been reported in which each antibody molecule is suggested to engage multiple epitopes on a single receptor molecule (intramolecular receptor engagement; Henry et al., 2021). These biepitopic antibodies were found to possess enhanced affinity relative to their monoepitopic counterparts. These types of antibodies may be particularly useful for activating receptors that do not require clustering. Finally, given the potential liabilities of these complex (non-conventional) antibody formats, future investigations should thoroughly characterize the influence of the formats of biepitopic antibodies on their biophysical properties and *in vivo* pharmacokinetics.

Previous reports indicate the promise of biepitopic antibodies for broad therapeutic applications (Bracken et al., 2021; Chi et al., 2022; Dai et al., 2021; DaSilva et al., 2020; Li et al., 2016; Tamaskovic et al., 2016; Xu et al., 2019). For example, biepitopic antibodies have recently been pursued in chimeric antigen receptor T cell (CAR-T) therapy for the treatment of B cell malignancies (Dai et al., 2021; Xu et al., 2019). Notably, the *in vivo* anti-tumor activity of CAR-T cells endowed with CD5 biepitopic targeting was much more effective relative to those that used monoepitopic targeting (Dai et al., 2021). Beyond applications relating to cellular activation, biepitopic targeting has been pursued for many applications involving inhibition of cellular processes, including for treating cancer and infectious disease (Bracken et al., 2021; Chi et al., 2022). Previous work highlights the utility of biepitopic targeting of tyrosine kinase receptors for anti-cancer applications via multiple mechanisms (DaSilva et al., 2020; Li et al., 2016; Tamaskovic et al., 2016). For example, HER2-targeted biepitopic antibody-drug conjugates (ADCs) showed enhanced ADC internalization and improved bioactivity relative to their monoepitopic counterparts (Li et al., 2016). Biepitopic antibodies have also shown promise for controlling receptor trafficking (DaSilva et al., 2020) and overcoming compensatory pro-mitogenic signaling in cancer models (Tamaskovic et al., 2016). We expect many more creative and impactful uses of biepitopic antibodies in the future, which can be generated in a simple and systematic manner using our competition-based discovery method.

### Significance

Agonist antibodies that target the TNF receptor superfamily enhance a wide range of immune functions. However, their therapeutic use faces multiple challenges, including their lack of intrinsic ability to efficiently cluster receptors and mediate strong activation without the need for secondary antibody clustering via other immune cells. Biepitopic antibodies that target two distinct epitopes on the same receptor possess an intrinsic ability to potently activate TNF receptors via receptor superclustering, but their generation requires laborious trial-and-error methods. Here, we report a facile method for selecting antibodies with unique pairs of epitopes and demonstrate how they can be combined to develop biepitopic antibodies that strongly activate multiple clinically important TNF receptors.

## Acknowledgements

We thank members of the Tessier lab for their helpful suggestions. This work was supported by the National Institutes of Health (R35GM136300, RF1AG059723, and R01AG080016 to P.M.T. and F32GM137513 to J.S.S.), National Science Foundation (CBET 1813963, 1605266, and 1804313 to P.M.T.), and the Albert M. Mattocks Chair (to P.M.T).

## Materials and methods

### Antibody discovery

The human nonimmune library (library #1; Feldhaus et al., 2003) and the synthetic library (library #2; Kelly et al., 2018) were expressed as Aga2-scFv fusion proteins on the surface of yeast using a standard yeast plasmid (pCTCON2). Library #1 was used for OX40 and library #2 was used for CD137. For library sorting, two initial MACS selections were conducted using human OX40-Fc or CD137-Fc (produced in-house). The antibody libraries were induced in SDGCAA media (2 g/L dextrose, 6.7 g/L yeast nitrogen base without amino acids, 5 g/L casamino acids, 20 g/L galactose, 8.56 g/L sodium phosphate monobasic monohydrate, 6.76 g/L sodium phosphate dibasic dihydrate) for 1 day at 30 °C to promote the expression of scFvs on the cell surface. 10^9^ yeast cells were incubated with 300 nM antigen (OX40-Fc or CD137-Fc) in 5 mL of 1X PBS supplemented with 1 g/L BSA (PBSB) and 1% milk at room temperature for 3 h. Post-incubation, the yeast cells were washed once with PBSB and incubated with 750 μL of Protein A microbeads (Miltenyi Biotec, 130071001) per 10^9^ cells, then gently rocked for 30 min at 4 °C. Next, the cells were rinsed with PBSB and passed through a MACS column to isolate cells bound to beads under a magnetic field. The captured beads were washed once with PBSB under the magnetic field and subsequently eluted into 5 mL of SDCAA media (20 g/L dextrose, 6.7 g/L yeast nitrogen base without amino acids, 5 g/L casamino acids, 16.75 g/L sodium citrate trihydrate, 4 g/L citric acid) at 30 °C. Subsequently, the cells were grown in SDCAA media for 1 day at 30 °C. Next, the enriched library was sorted via FACS at antigen concentrations of 300 nM for FACS rounds 1-3 against biotinylated OX40 ectodomain (Acro Biosystems, TN4-H82E4) and rounds 1-4 against biotinylated CD137 ectodomain (Acro Biosystems, 41B-H82E6). Following a 3 h incubation (as described above), yeast cells (10-fold excess number of cells as the remaining library diversity) were washed with PBSB and labeled with streptavidin-Alexa 647 (Jackson Laboratories, 016-600-084) at 1:1000 dilution for OX40 or neutravidin-PE (Invitrogen, A2660) at 1:300 for CD137 for antigen binding, and mouse anti-myc tag (primary antibody; Cell Signaling, 2276S) at 1:1000 dilution and goat anti-mouse IgG AlexaFluor 488 (secondary antibody; Invitrogen, A11001) at 1:200 dilution for antibody display. For FACS rounds 4-5 (OX40) and rounds 5-6 (CD137), 11D4 IgG or utomilumab IgG were prepared at equimolar concentrations relative to the receptor ectodomain for the first round of competition sorting, and 10-fold excess for the terminal round of sorting. Finally, the cells were incubated with the ectodomain/IgG complexes for 3 h at room temperature on a rocker prior to labeling with streptavidin-Alexa 647 or neutravidin-PE and mouse anti-myc tag and goat anti-mouse IgG AlexaFluor 488, as described above. Finally, the library was sorted using FACS and collected into 2 mL of SDCAA media. The cells were finally regrown and sequenced.

### Competitive binding analysis

Yeast cells expressing OX40 single-chain antibodies were prepared at 10^5^ cells per well in a 96-well plate and washed twice with PBSB. They were then combined with different concentrations of biotinylated human OX40 ectodomain (Acro Biosystems, TN4-H82E4) with or without a six-fold molar excess of soluble 11D4 IgG in PBSB containing 1% milk. Afterward, mouse anti-myc tag antibody (Cell Signaling, 2276S) was added at 1:1000 dilution for 3 h at room temperature under gentle agitation. Following the antigen incubation step, yeast cells were centrifuged at 2500 xg for 5 min, washed once with ice-cold PBSB, and incubated with labeling reagents. These reagents were goat anti-mouse IgG AlexaFluor 488 (Invitrogen, A11001) at 1:200 dilution and streptavidin-Alexa647 (Jackson Laboratories, 016-600-084) at 1:1000 dilution, which were added on ice for 5 min. Afterward, the cells were washed once with ice-cold PBSB and analyzed by flow cytometry.

For the bead-based competition assay, antibodies were immobilized on Protein A magnetic beads (Pierce, 88846) following the manufacturer’s protocol. The beads were washed twice with PBSB, blocked with 10% milk in PBSB, mixed end-over-end at room temperature for 1 h, and washed once more with PBSB. In a 96-well plate, the beads (10^5^ beads/well) were incubated with 100 nM biotinylated OX40 extracellular domain (Acro Biosystems, TN4-H82E4) in the presence or absence of a competition antibody (five-fold molar excess) in PBSB with 1% milk at room temperature for 3 h. Following the incubation, the beads were washed once with ice-cold PBSB and then incubated with streptavidin-Alexa647 (Jackson Laboratories, 016-600-084) at 1:1000 dilution on ice for 5 min. After labeling, the beads were washed once with ice-cold PBSB and evaluated by flow cytometry.

### Isolation and sequencing of single-chain antibodies

The terminal sorts of the yeast-displayed libraries were miniprepped using the Zymoprep Yeast Plasmid Miniprep Kit (Zymo Research, D2004) to recover enriched yeast plasmids. To recover the scFv library genes, plasmids were first transformed into DH5α bacterial cells and plated overnight at 37 °C on Luria Broth (LB) agar plates with 100 μg/mL ampicillin. After incubation, individual bacterial colonies were picked and grown overnight at 37 °C in LB media supplemented with ampicillin, and then miniprepped using the QIAprep Spin Miniprep Kit (Qiagen, 27106) for sequencing.

### Soluble antibody expression and purification

Following scFv sequence identification, gene fragments (gBlocks, Integrated DNA Technology) were designed to incorporate enriched scFv sequences at the C-termini of the light chains of the IgGs. To express these tetravalent antibodies, HEK 293-6E cells were grown and passaged in F17 media (Fisher Scientific, A1383502) supplemented with glutamine (Invitrogen, 2530081), Kollipher (Fisher Scientific, NC0917244), and G418 (Gibco, 10131027) at 1.5-2 million cells/mL. Next, 15 μg of vector plasmid (7.5 μg of variable light chain (V_L_) plasmid and 7.5 μg of variable heavy chain (V_H_) plasmid for bivalent and 11.25 μg of V_L_ plasmid and 3.75 μg of V_H_ plasmid for tetravalent antibodies) and PEI (3-fold excess, 45 μg) were mixed in F17 media for 10-15 min at room temperature then added to the cells. 24-48 h after transfection, cells were fed using 20% Yeastolate (Gibco, 292804) followed by another 2-5 day growth period at 37 °C. After protein expression, cells were collected via centrifugation at 2500 xg for 45 min and transferred to new tubes. Next, 0.5-1 mL of Protein A beads (Thermo Fisher Scientific, 89898) were added and gently rocked overnight at 4 °C. Protein A beads were collected by vacuum filtration and washed with 50-100 mL of 1X PBS. Protein A beads were then incubated in glycine buffer (0.1 M, pH 3) for 15 min at room temperature to elute the protein. Then, the eluted protein was buffer exchanged once into acetate buffer (20 mM, pH 5) using Zeba Spin Desalting Columns (Thermo Fisher Scientific, PI89890), after which the protein was filtered with 0.2 μm filters and the concentration was determined using a NanoDrop to measure protein A280. Finally, protein purity was analyzed by size-exclusion chromatography (SEC) and SDS-PAGE, followed by an additional purification step by SEC.

### OX40 and CD137 HEK-293T binding analysis

HEK-293T cell lines were first engineered to stably express doxycycline-inducible human OX40 or CD137 receptors (developed in-house) using a lentiviral delivery system. In 96-well plates, 50,000 OX40 or CD137 cells were plated with 100 ng/mL doxycycline to a volume of 150 μL, then incubated at 37 °C with 5% CO_2_ for 2 days. Then, antibodies were added to a final volume of 200 μL and incubated on ice for 1 h. Next, cells were centrifuged at 300 xg for 3 min and washed with ice-cold 1X PBS. After centrifugation, the wells were incubated with Alexa647 goat anti-human IgG Fcγ fragment antibody (Jackson ImmunoResearch Laboratories, 109-605-098) at 1:300 dilution for 4 min on ice. Cells were then centrifuged and washed with 150 μL of 1X PBS, as described above, then analyzed by flow cytometry.

### OX40 Jurkat T cell assay

The OX40 Bioassay (Promega, JA2191) was conducted following the manufacturer’s protocol. Briefly, thaw-and-use NFkB-*luc2*/OX40 Jurkat cells were seeded at a density of 50,000 cells/well in 96-well plates (Costar, 3917) and incubated at 37 °C with 5% CO_2_ overnight. Serial-diluted antibodies or controls were added and incubated for an additional 5 hours at 37 °C with 5% CO_2_. The assay plates were then equilibrated to room temperature for 10 min and 80 μL of Bio-Glo Reagent was added to each well. The assay plates were then incubated at room temperature for 5 min and luminescence was measured using the SpectraMax M3 microplate reader (Molecular Devices) set at 500 ms integration/well.

### CD4^+^ T cell proliferation assay

Human peripheral blood CD4^+^ T cells (Stem Cell Technologies, 200-0165) were cultured in ImmunoCult-XF T Cell Expansion media (Stem Cell Technologies, 10981) at 37 °C with 5% CO_2_. For the experimental assay, T cells were stained with carboxifluorescein diacetate succinimidyl ester (CFSE) using the CellTrace CFSE Cell Proliferation Kit (Invitrogen, C34554) following the manufacturer’s protocol. Briefly, cells were resuspended in 5 mL of serum-free 1X PBS supplemented with CFSE dye (5 μM working concentration) and incubated at room temperature for 20 min. Then, 25 mL of media with 10% FBS was added and incubated at room temperature for 5 min. Cells were then centrifuged at 270 xg for 4 min and resuspended in serum-free media. CFSE-labeled cells were seeded at a density of 50,000 cells/well in a 96-well plate. Next, serial-diluted antibodies were added over a range of concentrations, and ImmunoCult Human CD3/CD28 T Cell Activator (Stem Cell Technologies, 10971) was added at 5 μL/mL to a total volume to 200 μL. After incubating at 37 °C with 5% CO_2_ for 6 days, the cells were stained with BV510 anti-human CD4 (BioLegend, 357420) at 1:200 dilution on ice for 5 min. Cells were then centrifuged at 300 xg for 4 min, washed with 1X PBS, and analyzed via flow cytometry. Specifically, FlowJo v10.9 Software (BD Life Sciences) was used to analyze cell proliferation and calculate division indices (the average number of cell divisions for a T cell in the original population) with the FlowJo Proliferation Platform tool.

### CD8^+^ T cell proliferation assay

Human peripheral blood CD8^+^ T cells (Stem Cell Technologies, 200-0164) were cultured in ImmunoCult XF T Cell Expansion media (Stem Cell Technologies, 10981) at 37 °C with 5% CO_2_. Antibodies were immobilized on Dynabeads M-280 Tosylactivated (Invitrogen, 14203) as described previously (Desai et al., 2022). Briefly, a total of 10 μg of antibody consisting of 2 μg of anti-human CD3 antibody (Bio X Cell, BE0001-2) and 8 μg of experimental antibody in 1 mL of 1X PBS was incubated at room temperature for 2 days with rotation. The beads were subsequently blocked with 10 mM glycine buffer for 1 h with rotation, followed by washing twice with PBSB. Next, the CD8^+^ T cells were plated at a density of 50,000 cells/well in a 96-well plate to a volume of 150 μL. Then, 50 μL of bead solution was added at a ratio of 4:1 or 8:1 of conjugated beads:T cells. After incubating at 37 °C with 5% CO_2_ for 96 h, the cells were stained with BV510 anti-human CD8 (BioLegend, 344732) and APC anti-human CD25 (BioLegend, 302610) antibodies at 1:200 dilution each on ice for 5 min. Cells were then centrifuged at 300 xg for 4 min, washed with 1X PBS, and analyzed via flow cytometry.

