## Supplemental Figures for "Facile generation of biepitopic antibodies with intrinsic agonism for activating receptors in the tumor necrosis factor superfamily"

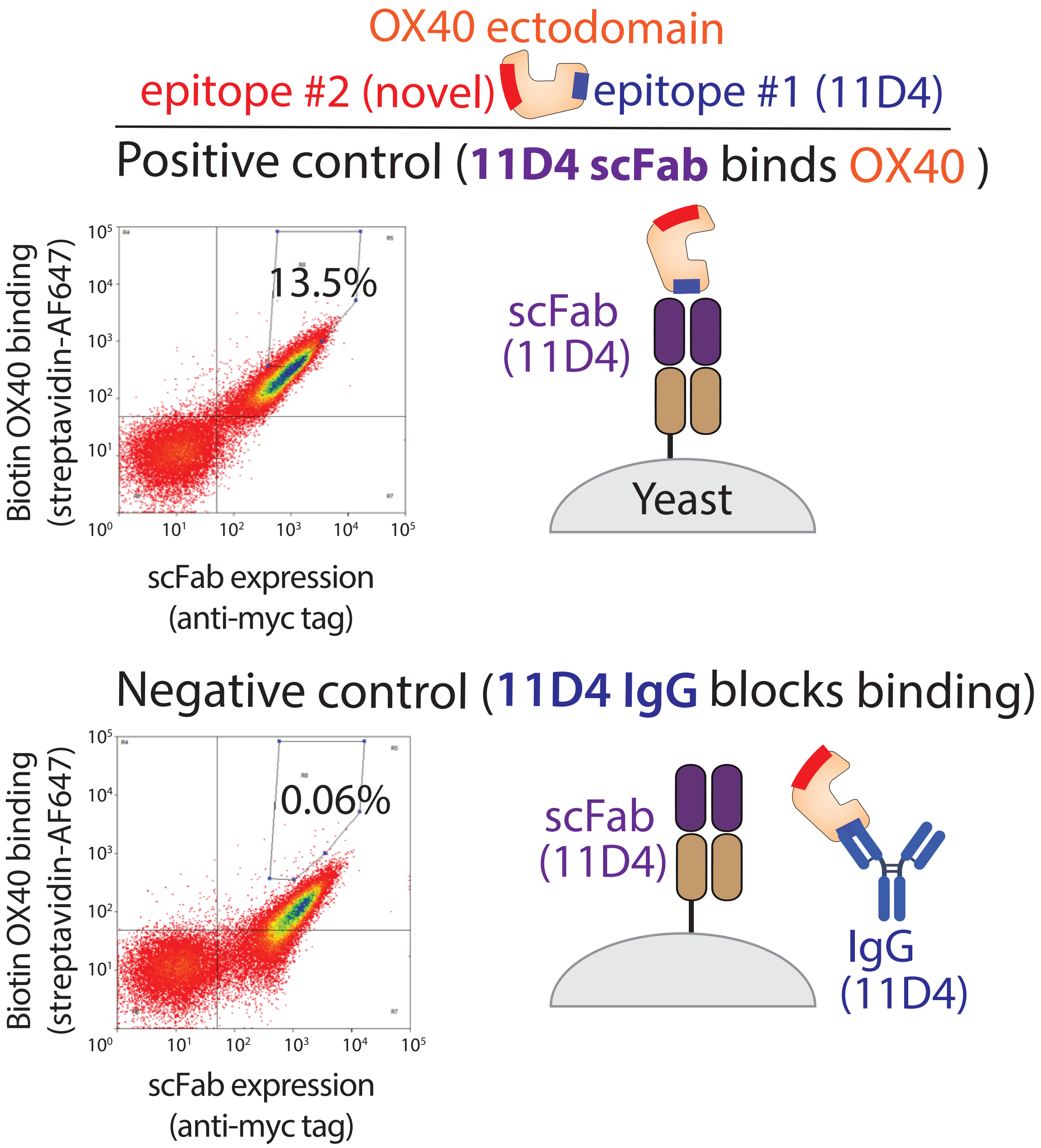


**Figure S1. Flow cytometry analysis of the 11D4 single-chain antibody displayed on yeast binding to human OX40 in the absence and presence of 11D4 IgG.** (Top) The parental 11D4 IgG was reformatted as a single-chain Fab (scFab), displayed on yeast, and confirmed to bind strongly to the OX40 ectodomain. (Bottom) The OX40 ectodomain was pre-blocked with 11D4 IgG and confirmed to bind weakly to the OX40 scFab on yeast due to epitope blocking.

A

```
OX.F1
QVQLQQSGPGLVKPSQTLSLTCAISGDSVSSNSVSWDWIRQSPSRGLEWLGRTYYRSKWYNEYAVSVESRI
TINPDTSKNQFSLQLNSVTPEDTAIYFCVRNNYFFDLWGRGTLVTVSSGILGSGGGGSGGGGSGGGGSEIV
LTQSPATLSLSPGERATLSCRASQSVSSYLAWYQQKPGQAPRLLIYDASDRATGIPARFSGSGSGTDFTLT
ISSLEPEDFAVYYCQLRSNWPPGYTFGQGTKVEIK

OX.F2
QVQLQQSGPGLVKPSQTLSLTCGISGDSVSSNSVSWDWIRQSPSRGLEWLGRTYYRSKWYNEYAVSVESRI
TINPDTSKNQFSLQLNSVTPEDTAIYFCVRNNYFFDLWGRGTLVTVSSGILGSGGGGSGGGGSGGGGSEIV
LTQSPATLSLSPGERATLSCRASQSVSSYLAWYQQKPGQAPRLLIYDASNRATGIPARFSGSGSGTDFTLT
ISSLEPEDFAVYYCQQRSNWPPMYTFGQGTKLEIK

OX.F3
QVQLQQSGPGLVKPWQTLSLTCGISGDSVSSNSVSWDWIRQSPSRGLEWLGRTYYRSKWYNEYAVSVESRI
TINPDTSKNQFSLQLNSVTPEDTAIYFCVRNNYFFDLWGRGTLVTVSSGILGSGGGGSGGGGSGGGGSEIV
LTQSPATLSLSPGERATLSCRASQSVSSYLAWYQQKPGQAPRLLIYDASNRATGIPARFSGSGSGTDFTLT
ISSLEPEDFAVYYCQQRSNWPPMYTFGQGTKLEIK
```

B

```
OX.F1      QVQLQQSGPGLVKPSQTLSLTCAISGDSVSSNSVSWDWIRQSPSRGLEWLGRTYYRSKWY    60
OX.F2      QVQLQQSGPGLVKPSQTLSLTCGISGDSVSSNSVSWDWIRQSPSRGLEWLGRTYYRSKWY    60
OX.F3      QVQLQQSGPGLVKPWQTLSLTCGISGDSVSSNSVSWDWIRQSPSRGLEWLGRTYYRSKWY    60
           ************** *******.*************************************

OX.F1      NEYAVSVESRITINPDTSKNQFSLQLNSVTPEDTAIYFCVRNNYFFDLWGRGTLVTVSSG    120
OX.F2      NEYAVSVESRITINPDTSKNQFSLQLNSVTPEDTAIYFCVRNNYFFDLWGRGTLVTVSSG    120
OX.F3      NEYAVSVESRITINPDTSKNQFSLQLNSVTPEDTAIYFCVRNNYFFDLWGRGTLVTVSSG    120
           ************************************************************

OX.F1      ILGSGGGGSGGGGSGGGGSEIVLTQSPATLSLSPGERATLSCRASQSVSSYLAWYQQKPG    180
OX.F2      ILGSGGGGSGGGGSGGGGSEIVLTQSPATLSLSPGERATLSCRASQSVSSYLAWYQQKPG    180
OX.F3      ILGSGGGGSGGGGSGGGGSEIVLTQSPATLSLSPGERATLSCRASQSVSSYLAWYQQKPG    180
           ************************************************************

OX.F1      QAPRLLIYDASDRATGIPARFSGSGSGTDFTLTISSLEPEDFAVYYCQLRSNWPPGYTFG    240
OX.F2      QAPRLLIYDASNRATGIPARFSGSGSGTDFTLTISSLEPEDFAVYYCQQRSNWPPMYTFG    240
OX.F3      QAPRLLIYDASNRATGIPARFSGSGSGTDFTLTISSLEPEDFAVYYCQQRSNWPPMYTFG    240
           ***********:************************************ ****** ****

OX.F1      QGTKVEIK   248
OX.F2      QGTKLEIK   248
OX.F3      QGTKLEIK   248
           ****:***
```

**Figure S2. Amino acid sequences of single-chain OX40 antibodies.** (A) The scFv amino acid sequences are presented in the following format: variable heavy region - linker region - variable light chain region. (B) Multiple amino acid sequence alignment performed using Clustal Omega Sequence Alignment tool. "*" indicates perfect alignment; ":" indicates similar residues; "." indicates weakly similar residues. Heavy chain CDRs 1-3 (blue), linker region (grey), light chain CDRs 1-3 (green), and mutations (red) are shown. CDR residues were determined using Kabat numbering.

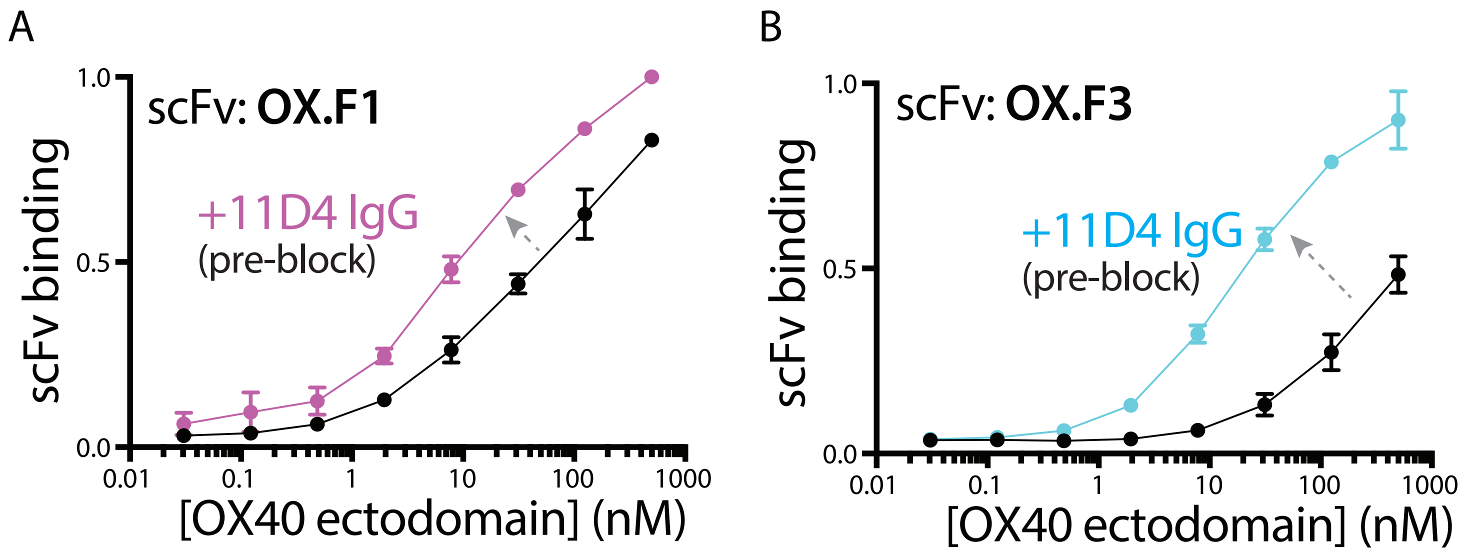


**Figure S3. Selected single-chain antibodies possess unique OX40 epitopes relative to 11D4 IgG.** The candidate single-chain antibodies, including (A) OX.F1 and (B) OX.F3, were displayed on yeast and demonstrated to bind OX40 ectodomain even when the 11D4 epitope is pre-blocked. The results are averages of three independent experiments and the error bars are standard deviations.

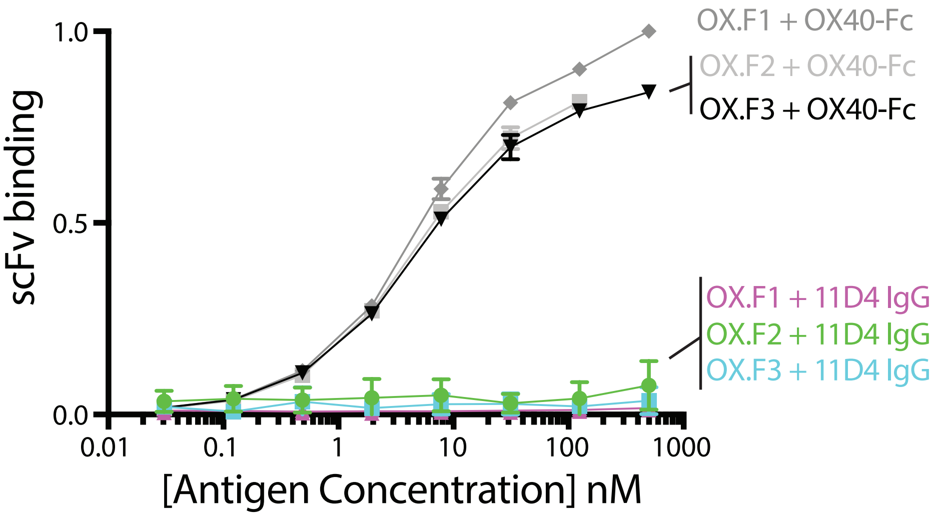


**Figure S4. OX40 single-chain antibodies lack affinity for 11D4 IgG.** Selected OX40 scFvs were displayed on the surface of yeast and tested for binding to 11D4 IgG using flow cytometry. As a control, the OX40 scFvs were confirmed to bind OX40 (as an Fc fusion protein). The results are averages of two independent experiments and the error bars are standard deviations.

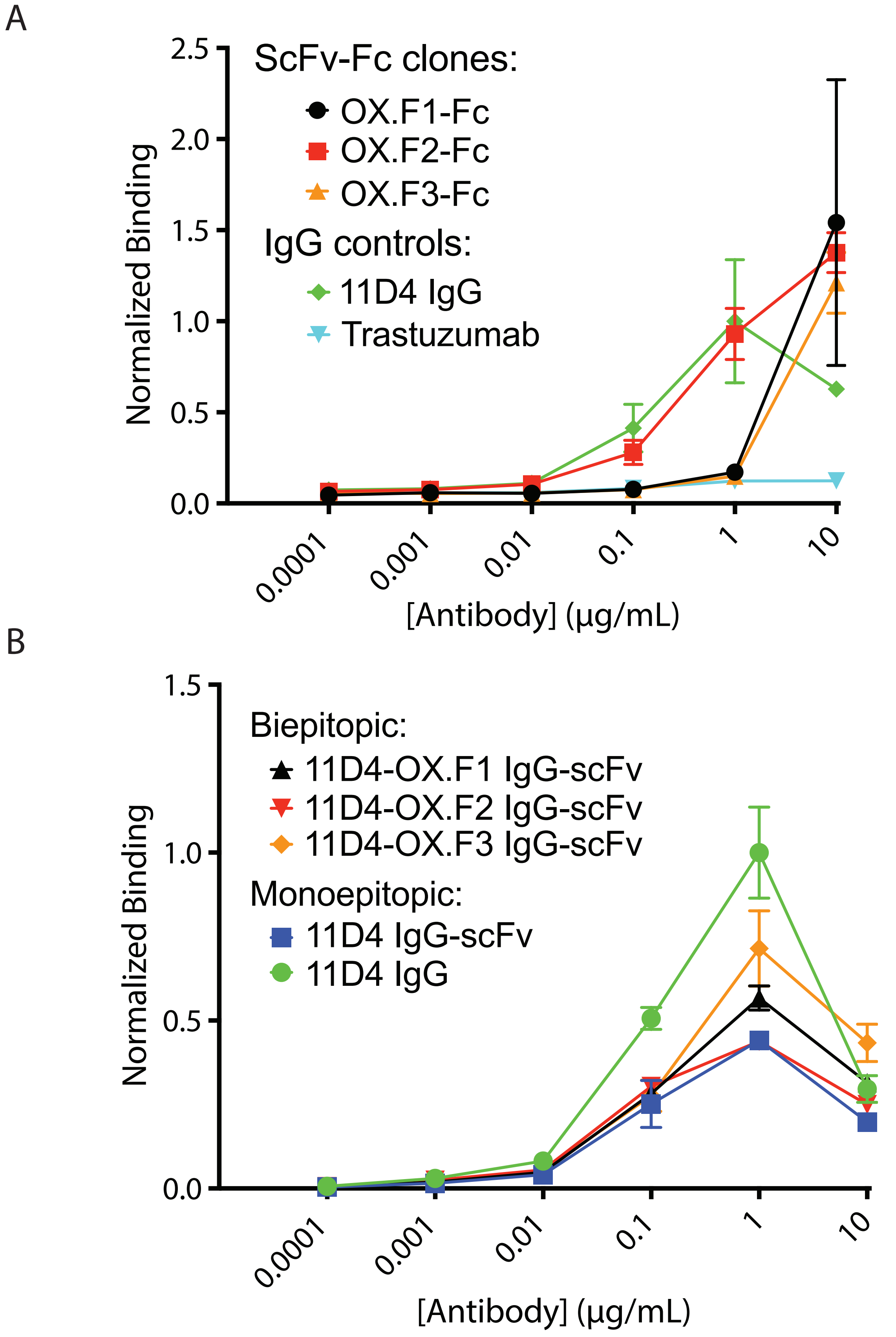


**Figure S5. scFv-Fc and IgG-scFv binding to OX40 on HEK293 cells.** (A) Selected scFvs were generated as scFv-Fc fusion proteins and their binding to OX40 on HEK293 cells was evaluated. (B) The binding of biepitopic (tetravalent) and monoepitopic (tetravalent and bivalent) antibodies to OX40 on HEK293 cells was evaluated. The results are averages of three independent experiments and the error bars are standard deviations.

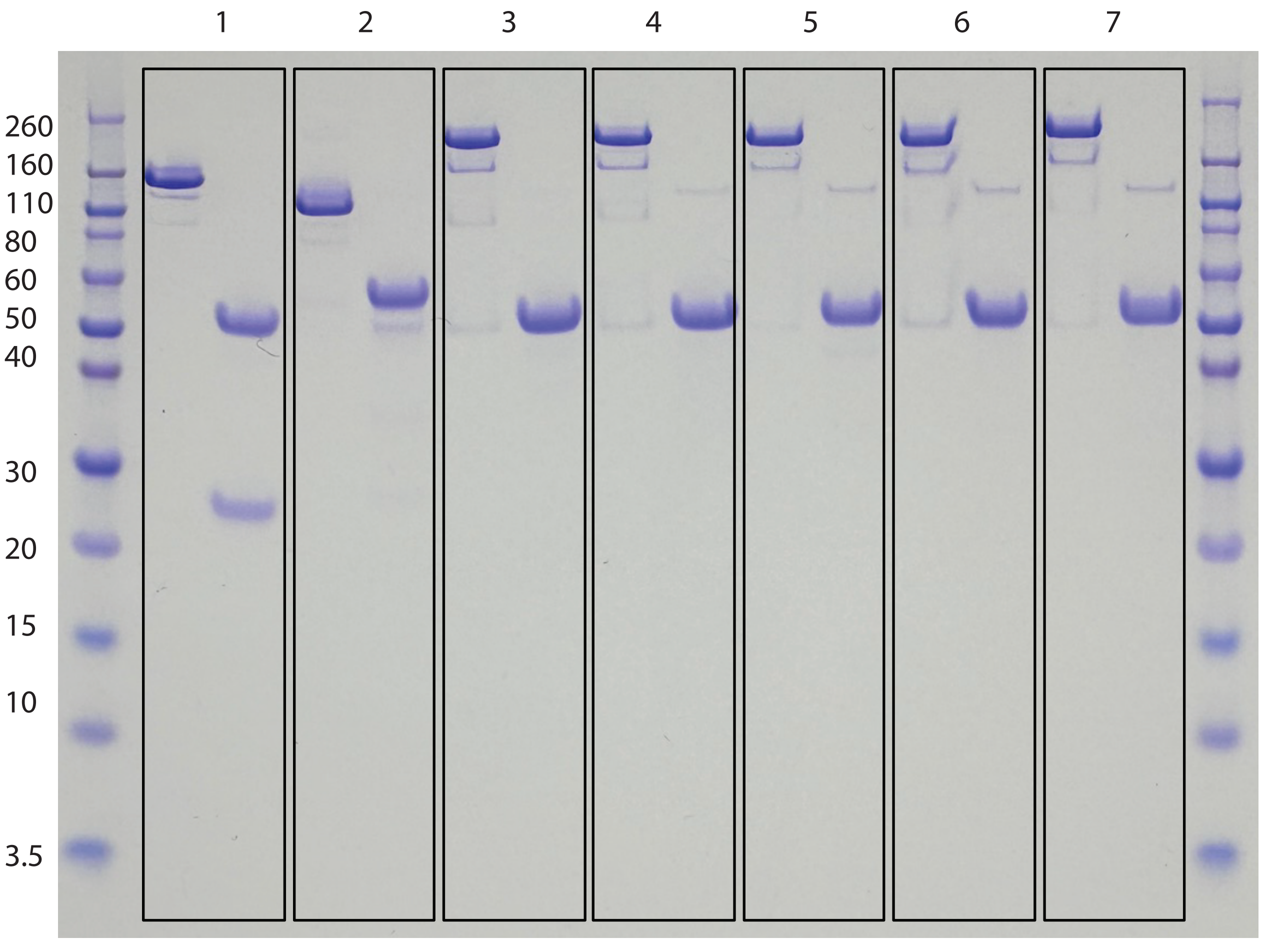


1: 11D4 IgG
2: OX.F2 scFv-Fc
3: 11D4 IgG-scFv
4: 11D4-OX.F2
5: tela-OX.F2
6: poga-OX.F2
7: 18D8-OX.F2

**Figure S6. SDS-PAGE analysis of OX40 monoepitopic and biepitopic antibodies.** The antibodies were evaluated in non-reducing (left lane) and reducing (right lane) conditions.

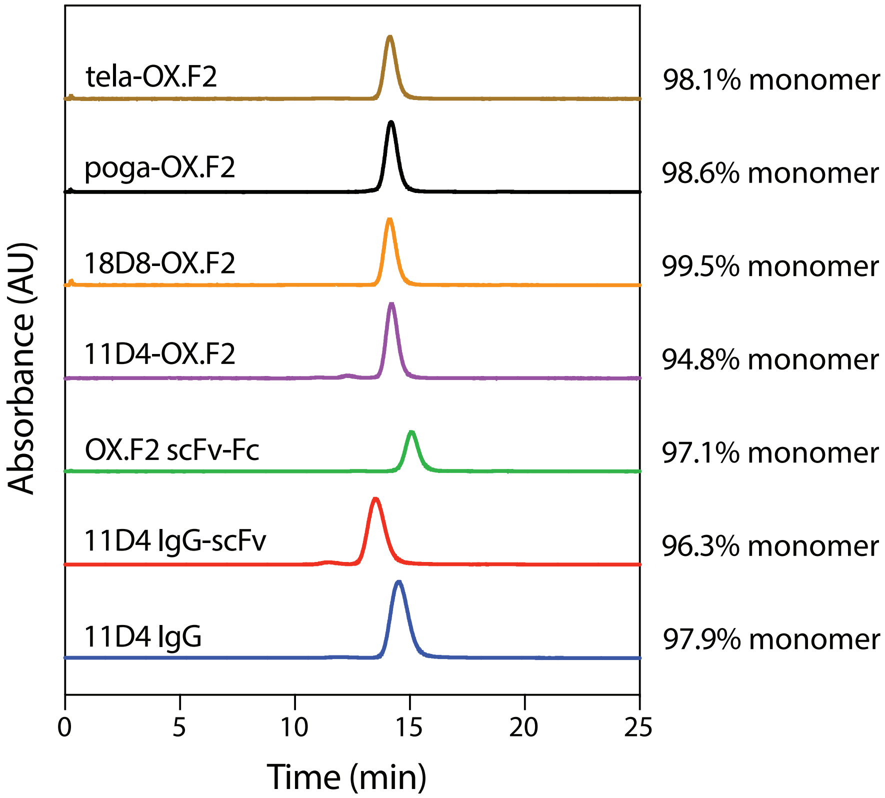


**Figure S7. Size-exclusion chromatography analysis of OX40 monoepitopic and biepitopic antibodies.** Antibody purity (percent monomer) was evaluated by size-exclusion chromatography.

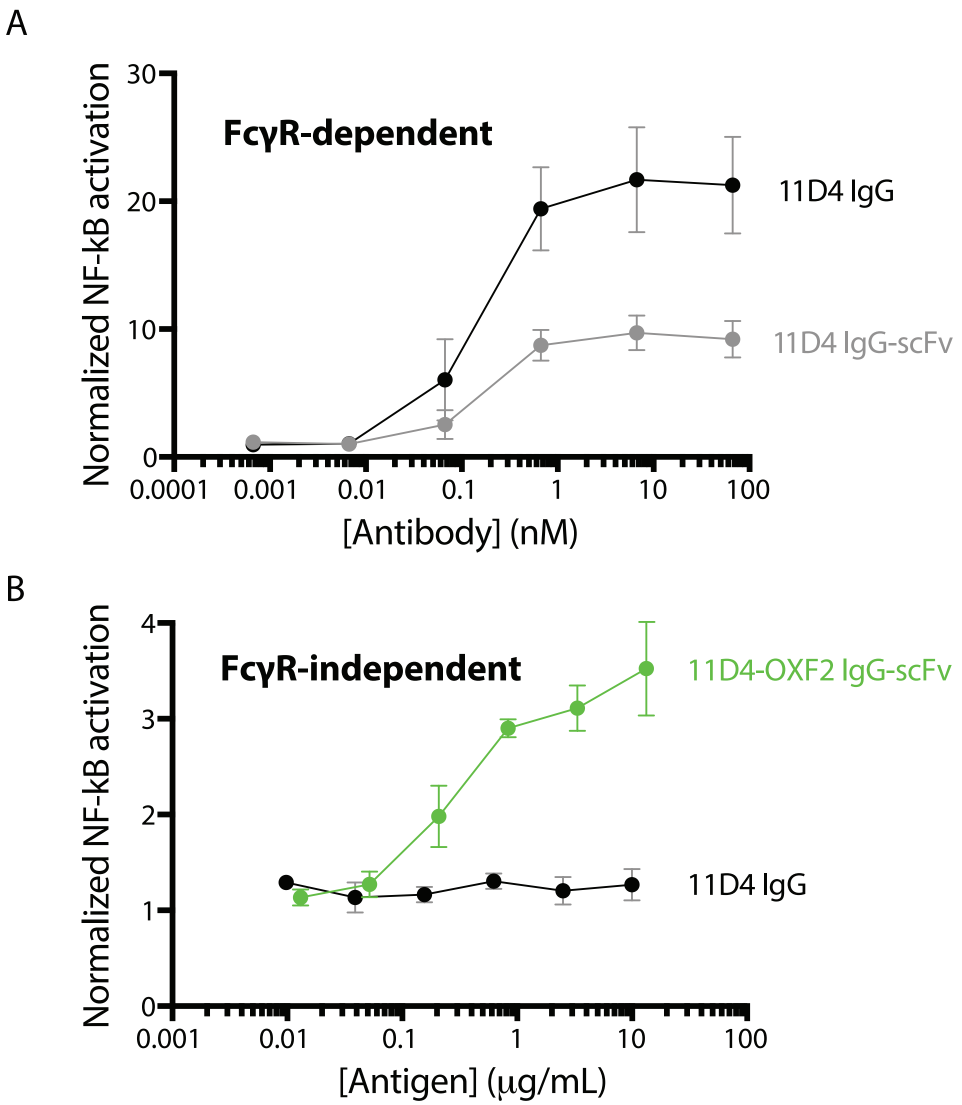


**Figure S8. Evaluation of FcγR-dependent and independent agonism of human OX40 using a Jurkat T cell assay.** The antibodies were evaluated for their ability to promote NF-κB activation in an OX40+ reporter Jurkat T cell line (A) with FcγR-dependent linking via CHO-K1 cells overexpressing FcγRIIb or (B) without Fc-mediated crosslinking. The results are averages of three or more independent experiments and the error bars are standard deviations.

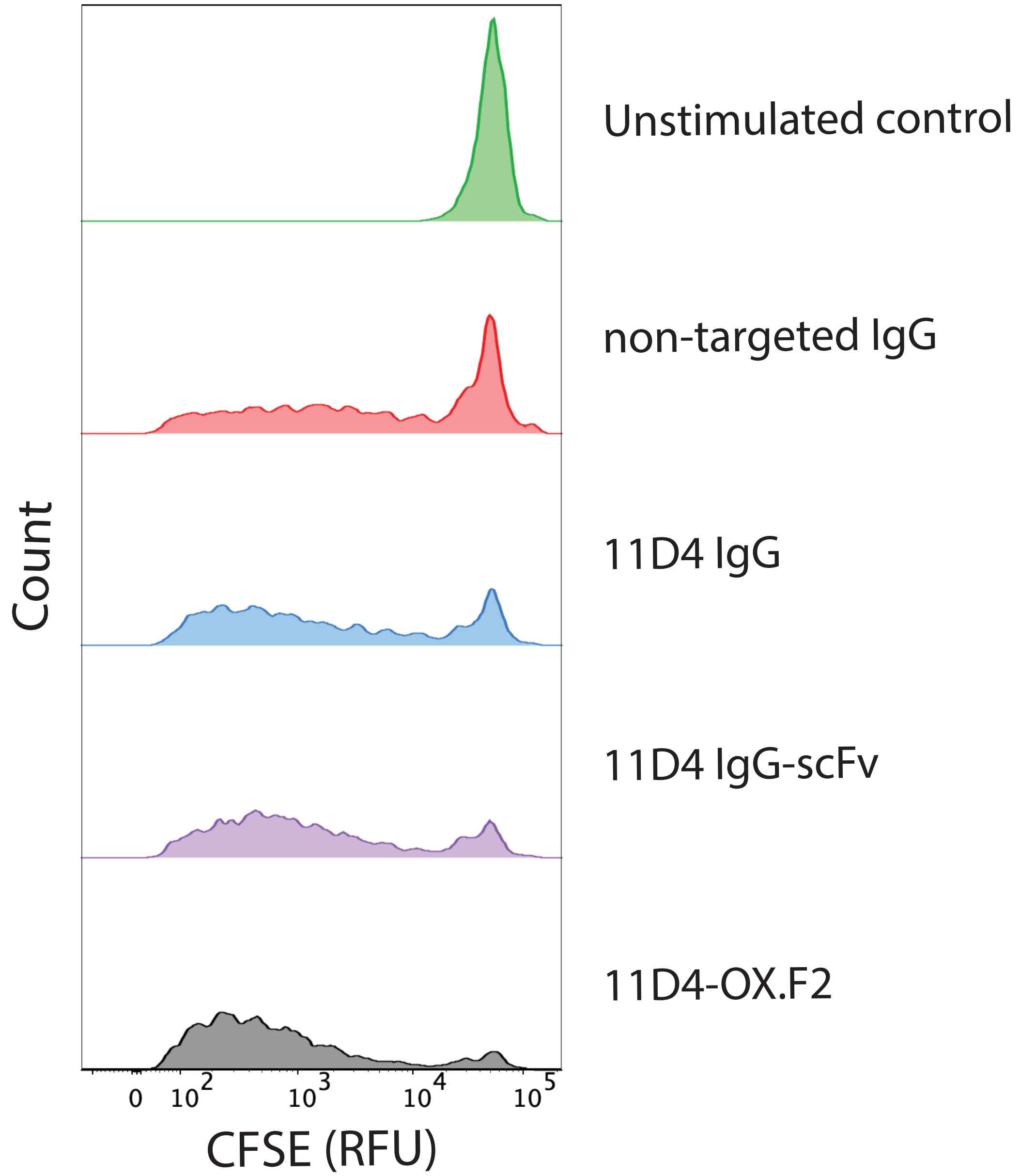


**Figure S9. Biepitopic OX40 antibody induces stronger human CD4+ T cell proliferation than monoepitopic OX40 antibodies.** Representative histograms of human CD4+ T cell proliferation as measured by CFSE expression following 6-day incubation with various antibodies and 5 µL/mL CD3/CD28 T cell activator solution. The biepitopic, tetravalent antibody (11D4-OX.F2) induced stronger proliferation than the monoepitopic, tetravalent antibody (11D4 IgG-scFv) and the monoepitopic, bivalent antibody (11D4 IgG).

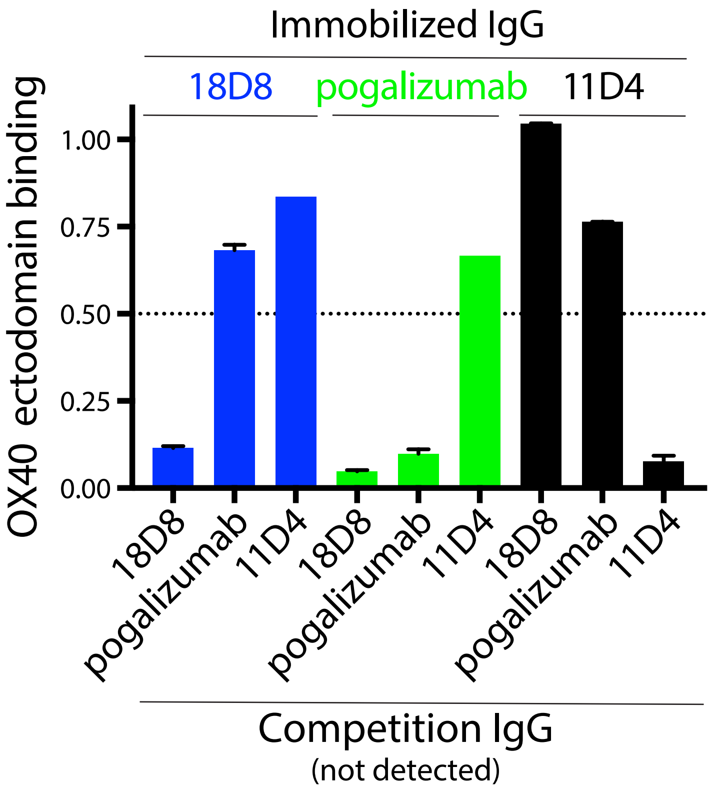


**Figure S10. Competitive binding analysis of OX40 IgGs.** Three OX40 IgGs were separately immobilized on Protein A beads, and their binding to OX40 was evaluated in the presence of different OX40 IgGs via flow cytometry. The results are averages of three independent experiments and the error bars are standard deviations.

```
CD.K2
DIQMTQSPSSLSASVGDRVTITCRASQSISSYLNWYQQKPGKAPKLLIYAASSLQSGVPSRFSGSGSGTDF
TLTISSLQPEDFATYYCQQSYSTPLTFGQGTKVEIKSGILGTTAASGSSGGSSSGAEVQLVQSGAEVKKPG
ESLKISCKGSGYSFTSYWIGWVRQMPGKGLEWMGIIYPGDSDTRYSPSFQGQVTISADKSISTAYLQWSSL
KASDTAVYYCARFTYYYYDYDYAGFDYWGQGTLVTVSS
```

**Figure S11. Amino acid sequence of the isolated CD137 single-chain antibody (CD.K2).** The scFv amino acid sequence is presented in the following format: variable light region - linker region - variable heavy chain region.

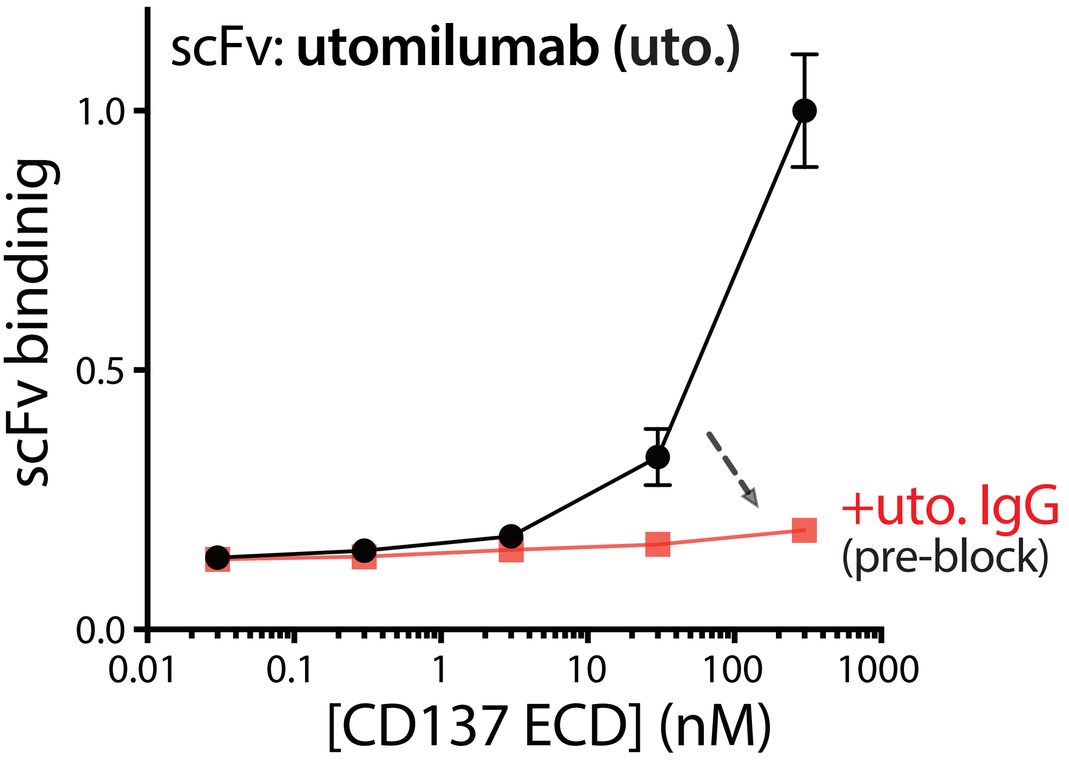


**Figure S12. CD137 binding to the utomilumab single-chain antibody on yeast is inhibited by utomilumab IgG.** The CD137 ectodomain was pre-incubated with uto. IgG (1:5 molar ratio) and then the binding of the CD137/u-to. complex to uto scFv was evaluated via flow cytometry.

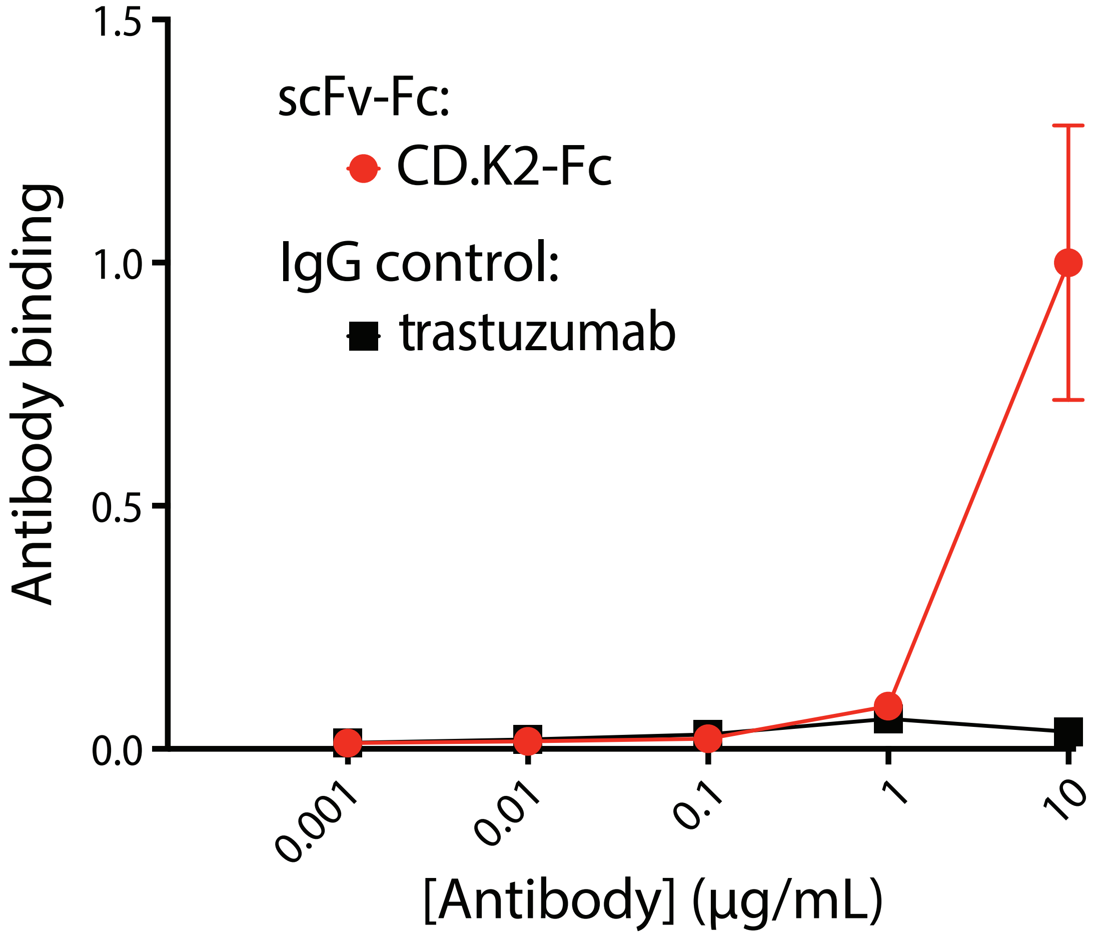


**Figure S13. scFv-Fc binding to CD137 on HEK293 cells.** (A) The selected scFv (CD.K2) was generated as an scFv-Fc fusion protein and its binding to CD137 on HEK293 cells was evaluated. The results are averages of median values and the error bars are standard deviations.

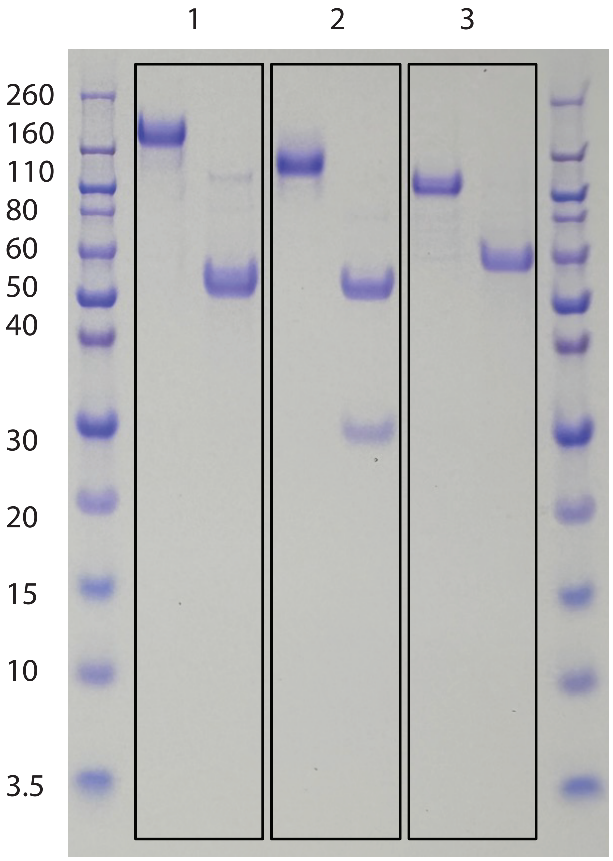


**Figure S14. SDS-PAGE analysis of CD137 antibodies.** The antibodies (1: uto-CD.K2; 2: uto. IgG; 3: CD.K2-Fc) were evaluated in non-reducing (left lane) and reducing (right lane) conditions.

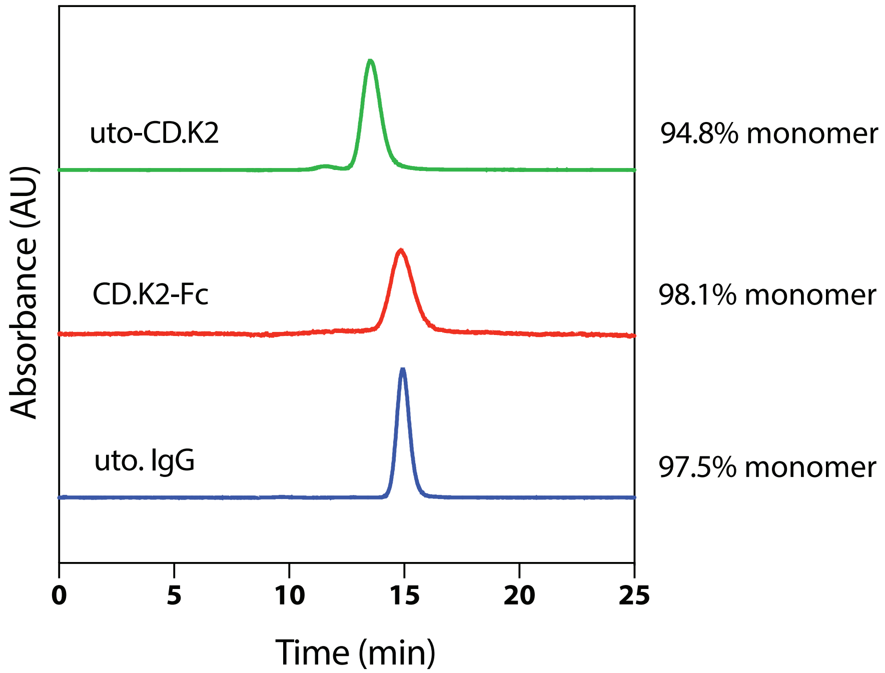


**Figure S15. Size-exclusion chromatography analysis of CD137 antibodies.** Antibody purity (percent monomer) was evaluated by size-exclusion chromatography.

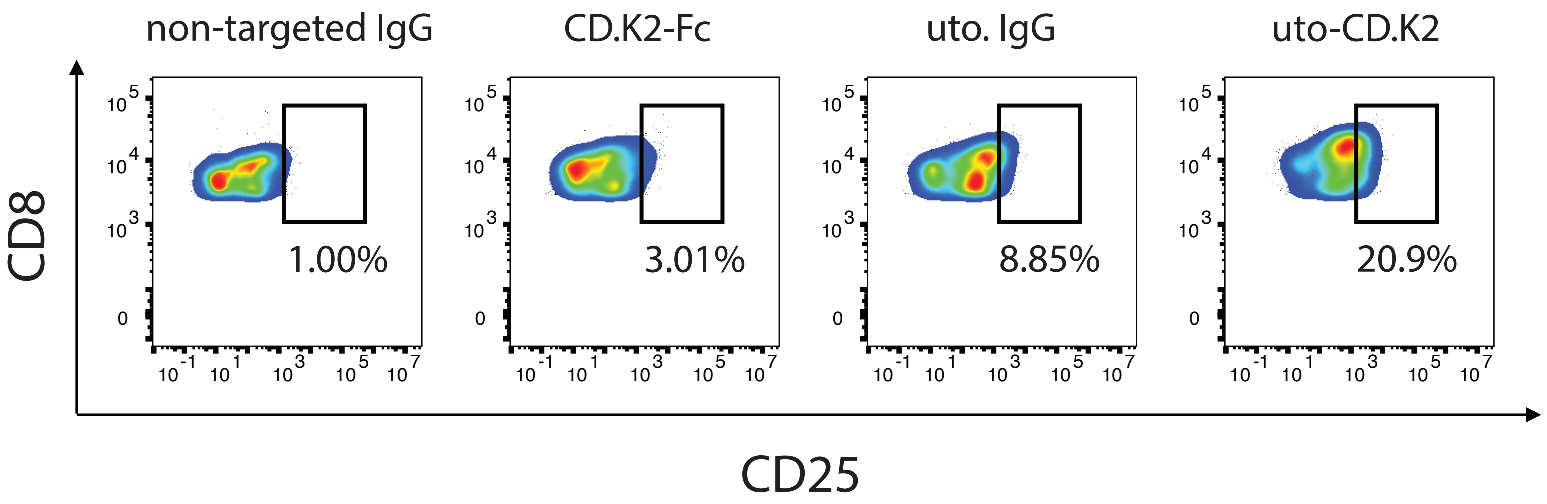


**Figure S16. Biepitopic CD137 antibody induces stronger human CD8+ T cell proliferation than monoepitopic CD137 antibody.** Representative FACS cytograms of human CD8+ T cell proliferation as measured by CD25 expression following 4-day incubation with various antibodies immobilized on beads. Data shown at 8:1 beads:T cells. The biepitopic antibody (uto-CD.K2) induced stronger CD25 expression relative to the monoepitopic antibodies (uto. IgG and CD.K2-Fc) and a control IgG.
